# Multivalent Afadin Interaction Promotes IDR-Mediated Condensate Formation and Junctional Separation of Epithelial Cells

**DOI:** 10.1101/2024.04.26.591237

**Authors:** Shuhei Kuno, Ryu Nakamura, Tetsuhisa Otani, Hideru Togashi

## Abstract

In epithelial cells, cell-cell adhesion is mediated by the apical junctional complex (AJC), which consists of tight junctions (TJs) and adherens junctions (AJs) aligned from the apical to the basal axis. However, the mechanism of AJC formation on the apical side and the separation of these junctions within AJCs are poorly understood. We found that multivalent interactions of afadin with adhesion molecules and the cytoskeleton lead to condensate formation in an intrinsically disordered region (IDR)-dependent manner, which promotes efficient accumulation in the linear AJ during initial junction formation. Furthermore, we found that afadin and ZO-1 were able to induce different condensate formations in the cell and that these molecules were differentially distributed from each other. These properties of afadin explain how it strictly localizes to AJs in epithelial cells and is involved in regulating the segregation of AJ and TJ within the AJC.

## Introduction

Epithelial cells form sheets and tubes constituting most organs and tissues in animals, and cell-cell adhesion plays a vital role in epithelial cell morphogenesis and function. The primary mediator of cell-cell adhesion is the apical junctional complex (AJC), which comprises tight junctions (TJs) and adherens junctions (AJs) that are typically aligned from the apical to basal sides^1–3^. In simple epithelia, cuboidal or columnar cells attach to each other through their lateral membranes. At the ultrastructural level, where the linear AJ is known as the zonula adherens (ZA) and is associated with a bundle of actin filaments, below the ZA, junctional structures extend to the basal ends of the cells, organizing the “lateral membrane contacts” (LCs)^4^. Although LCs span the majority of junctions, they are linked to less-organized actin filaments^5^. The mechanism by which the AJC forms on the apical side of the cell is not well understood, and the molecular mechanism by which TJs and AJs separate into apical and basal sides within the AJC remains unclear.

Previous studies have suggested that nectins initiate cell-cell adhesion and subsequently recruit the cadherin/catenin system to facilitate AJ formation^6^. Afadin, an anchoring protein that binds to nectin through the PDZ domain, localizes to AJs and interacts with various proteins, including cell adhesion molecules, signaling molecules, and the actin cytoskeleton^7,8^. Afadin has two splicing isoforms, l-afadin and s-afadin. Epithelial cells mainly express l-afadin, which is the longest isoform that binds to nectin and F-actin^9^, here simply referred to as afadin. Afadin-knockout (KO) mice exhibit impaired AJ formation and apical-basal polarity, as well as TJs in the neuroepithelium of the ectoderm, leading to embryonic lethality^10,11^. AJs are highly impaired in the ectoderm of afadin KO mice and embryoid bodies^10,12^. Additionally, mice lacking afadin in nephron precursors fail to develop normal apical-basal polarity, leading to diminished integration of nectin complexes and failure to recruit cadherin^13^. The phenotypes observed in afadin-KO mice suggest a crucial role of afadin in the formation of AJ and TJ. However, in cultured afadin-KO cells, there is evidence of cadherin-dependent cell adhesion and TJ formation, indicating phenotypic inconsistency with afadin-KO mice^14,15^. Canoe, the *Drosophila* homologue of afadin, was similarly identified as a component of AJ. A previous study showed that canoe is not essential for AJ assembly; however, its morphogenesis is impaired^13^. Although our understanding of the molecular interactions involving afadin and canoe has advanced, there is still a gap in our understanding of how afadin functions in AJ formation.

Afadin and nectins are strictly localized to AJs in epithelial cells, whereas cadherin/catenin complexes are distributed widely along LCs^9^. During the initial phases of junction formation and epithelial polarization, nectin/afadin and cadherin/catenin complexes localize to primordial AJs; however, their localization cannot be distinguished. Localization of nectin/afadin and cadherin/catenin complexes differs with epithelial polarity establishment^6^. However, the mechanism by which the nectin/afadin complex strictly localizes to the proper location of AJ in epithelial cells remains unknown. Recently, it has been reported that cells and tissues utilize a known domain- independent protein-autonomous assembly mechanism called liquid-liquid phase separation to maintain function, and the intrinsically disordered region (IDR)-dependent phase separation of ZO proteins drives TJ formation^16–18^. Despite having a long C- terminal IDR, as suggested by protein sequence predictions for canoe and afadin^19,20^, afadin is not predicted to be a phase-separating protein^21^. However, Gurley et al. recently reported severe morphological defects in head involution and dorsal closure in canoe mutants^19^. The severe phenotypes of the mutants were due to the loss of IDR, strongly suggesting the importance of an evolutionarily conserved IDR of canoe and afadin. This recent progress led us to hypothesize that IDR may play a role in the formation of AJ. In this study, we investigated the role of the C-terminal IDR of afadin during initial junction formation.

## Results

### Impairment of apical junction formation by loss of afadin

To investigate the role of afadin in the initial junction formation process in epithelial cells, we generated afadin-KO MDCK II (MDCK) cells, which possess typical epithelial junctions comprising AJCs and LCs. Successful KO was confirmed by western blotting^14^ **(Fig. S1B)**. Initially, we examined the localization pattern of afadin, α-catenin, and F-actin through immunofluorescence staining in wild-type (WT) cells 24 h following replating. Afadin signals were observed along the apical junctions, but hardly on the LCs **(Fig. 1A)**. F-actin exhibited prominent localization along the apical edges of the junctions in a linear cable fashion and displayed an amorphous pattern in the LCs **(Fig. 1A, Boxed regions)**. α-catenin signals primarily localized at the apical junctions and were also detected at LCs **(Fig. 1A)**. Signals for α-catenin and F-actin colocalized at apical junctions and LCs. In afadin-KO cells, the linear signals of α-catenin and F-actin at apical junctions were markedly reduced, while the areal signals on the lateral side were notably increased **(Fig. 1A, X-Z)**. These observations were confirmed by densitometric analysis **(Fig. 1A, graph)**. Loss of linear localization of α-catenin and F-actin at the apical junctions indicated an impairment in linear AJ formation due to afadin KO.

**Figure 1.**
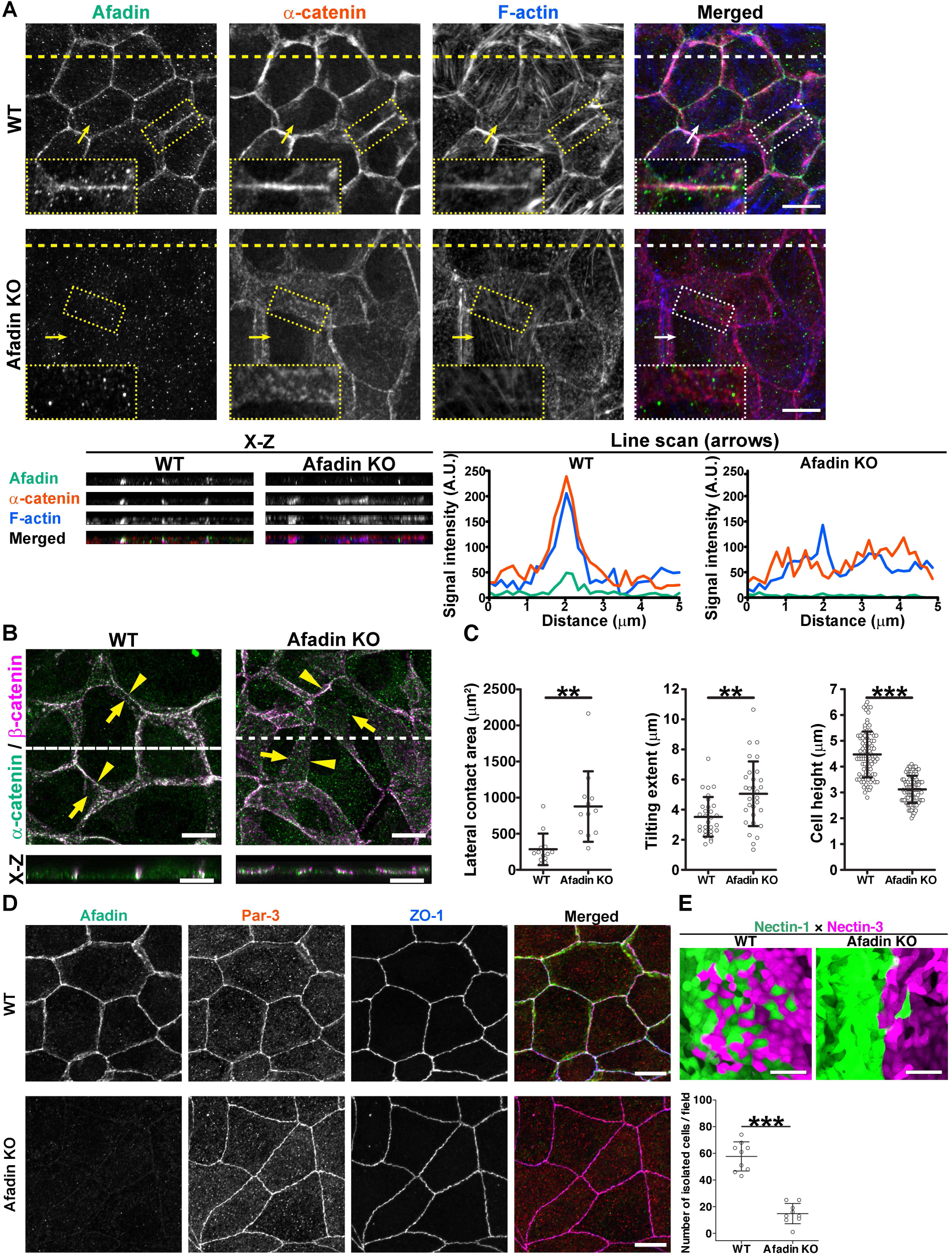
Impairment of apical junction formation by loss of afadin **(A)** Distribution of afadin (green), α-catenin (red), and F-actin (blue) in WT and afadin- KO cells; apical views of WT or afadin-KO MDCK cells cultured for 24 h and stained for the indicated molecules. Dashed lines indicate the position of vertical section. Arrows indicate scanned regions in the densitometric traces. A representative image is shown of six independent experiments. Scale bars, 10 μm. (X-Z) Vertical sections of the upper images. (Bottom, right) Line scans were performed along the arrows. A.U., arbitrary unit. **(B)** Distribution of α-catenin (green) and β-catenin (magenta) in WT and afadin-KO cells. Merged images are shown; apical views of WT or afadin KO cells cultured for 24 h and stained for α- and β-catenin. Arrowheads indicate the apical cell junctions. Arrows point to the basal edge of the junctions. Scale bars, 10 μm. (X-Z) Vertical sections of the upper images. Scale bars, 10 μm. **(C)** Quantifications of lateral contact area, tilting extent, and cell height. Whiskers represent mean ± standard deviation (S.D.). Lateral contact area; n = 13 for WT; n = 12 for afadin KO. Tilting extent; n = 30 for WT; n = 30 for afadin KO. Cell height; n = 95 for WT; n = 102 for afadin KO. Mann-Whitney *U*-test; **, P ≤ 0.01, ***, P ≤ 0.001. **(D)** Distribution of afadin (green), Par-3 (red), and ZO-1 (blue) in WT and afadin-KO cells. WT and afadin KO cells were cultured for 24 h and triple stained with the indicated molecules. Scale bars, 10 μm. **(E)** Mosaic patterning formation in EGFP and nectin-1 (green) or mScarlet and nectin-3 (magenta) expressing WT or afadin-KO MDCK cells. Scale bars, 50 μm. Whiskers represent the mean ± S.D. Fields, 397 μm^2^. n = 9 for EGFP and nectin-1; n = 9 for mScarlet and nectin-3. Mann-Whitney *U*-test; ***, P ≤ 0.001. See also STAR methods details. A representative image is shown of six independent experiments.

We also examined the localization of α- and β-catenins in WT and afadin-KO cells 24 h following replating. These catenins fully colocalized, respectively **(Fig. 1B)**. Simultaneously, the tilt extent and lateral contact area of LCs significantly increased in afadin-KO cells, accompanied by a reduction in cell height **(Fig. 1A,C)**. Despite these changes, the localization of the TJ-associated proteins, ZO-1 and Par-3, appeared nearly normal, with a sharp line along the apical edges. These observations suggested that TJ formation was minimally affected by afadin KO **(Fig. 1D)**. These results support the notion that afadin regulates the localization of linear AJs but does not affect TJ formation in cultured MDCK cells.

To clarify the functional significance of the apical AJ localization, we examined its impact on epithelial behavior using mosaic-forming assay^22^**(Fig. 1E)**. In the mosaic- forming assay, two different transfectants, which expressed EGFP or mCherry to distinguish them, were seeded into two wells that were separated by an insert. MDCK transfectants expressing nectin-1 and nectin-3, respectively, intermingled via cellular intercalations at the border; however, afadin-KO transfectants did not. These results indicated that afadin is required for the cellular intercalations in mosaic cellular patterning.

### Requirement of C-terminal IDR to apical localization of afadin

Afadin has multiple domains and is predicted to contain unique and extended IDRs in its C-terminal region^19,23^. To understand how afadin regulates apical junction formation in the epithelial cells, we conducted a domain analysis of afadin. Initially, we examined the junctional localization of EGFP-tagged afadin (EGFP-afadin) in afadin-KO cells **(Fig. 2A&S1A,C)**. EGFP-afadin was localized linearly along the apical junctions, fully restoring the localization of α-catenin and F-actin. We stably expressed various EGFP-afadin fragments in afadin-KO cells to understand the role of each domain in afadin localization. EGFP-afadinΔRA, EGFP-afadinΔPDZ, EGFP-afadinΔPR1-2, EGFP- afadinΔCC, and EGFP-s-afadin all localized linearly to apical junctions, similar to EGFP-afadin **(Fig. S1A,D)**. These results indicate that the function of a single domain cannot explain the apical localization of afadin.

**Figure 2.**
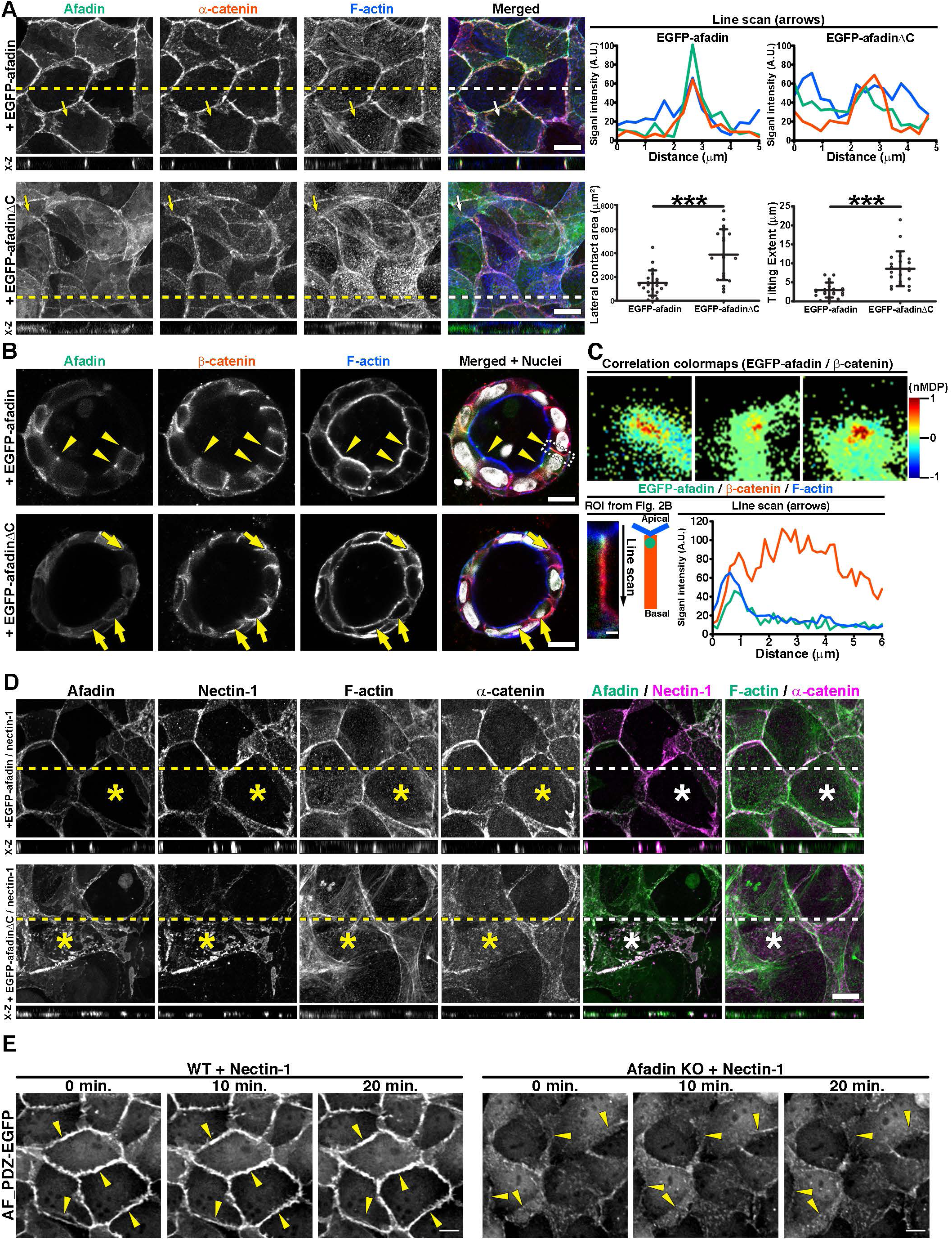
Destabilization of apical junction by lack of C-terminal region of afadin **(A)** Distribution of afadins (green), α-catenin (red), and F-actin (blue) in EGFP-afadin or EGFP-afadinΔC expressing afadin-KO cells. Apical views of the transfectants cultured for 24 h and stained for the indicated molecules. Dashed lines indicate the position of vertical section. Arrows indicate scanned regions in the line scans. Scale bars, 10 μm. (X-Z) Vertical sections of the upper images. (Upper, right) Line scan graphs were performed along the arrows. A.U., arbitrary unit. (Bottom, right) Quantification of lateral contact area and tilting extent of LCs. Whiskers represent the mean ± S.D. Lateral contact area; n = 20 for EGFP-afadin; n = 21 for EGFP-afadinΔC. Tilting extent; n = 20 for EGFP-afadin; n = 21 for EGFP-afadinΔC. Mann-Whitney *U*-test; ***, P ≤ 0.001. **(B)** Cyst formation of the cells expressing EGFP-afadin and EGFP-afadinΔC. These cysts were cultured for 8 days and stained for the indicated molecules. These images show the cross section of the cysts. Arrowheads indicate the apical cell junctions. Arrows point to the lateral contacts of the junctions. Scale bars, 10 μm. **(C)** Correlation colormaps of afadin and β-catenin of the cysts expressing EGFP-afadin. Colormaps indicate distribution of normalized mean deviation product of each pixel. A value of 1 (Red) reflects a positive correlation, while values close to -1 (Blue) indicate a negative correlation. nMDP, normalized mean deviation product. **(D)** Distribution of afadins, nectin-1, α-catenin, and F-actin in EGFP-afadin- or EGFP-afadinΔC- and nectin-1- expressing cells. Afadin-KO cells stably expressed EGFP-afadin or EGFP-afadinΔC and nectin-1 and were stained with the indicated molecules. Asterisks indicate cells of interest. Scale bars, 10 μm. (X-Z) Vertical sections of the upper images. A representative image is shown of six independent experiments. **(E)** Live-cell imaging of junctions in WT or afadin-KO cells expressing AF_PDZ-EGFP and nectin-1. Time-lapse images were taken to observe junction formation over a period of 20 min, with images captured at 10-min intervals. See also Video S1&S2. Scale bars, 10 µm. Arrowheads indicate the junctions of interest.

Next, we investigated the effect of the IDR-containing C-terminal region of afadin on its localization. We generated EGFP-afadinΔC, truncated of the IDR- containing C-terminal region but retaining the F-actin binding region, and stably expressed it in afadin-KO cells **(Fig. S1C)**. These afadin fragments and full-length afadin were stably introduced into afadin-KO cells. Low expression clones were selected by FACS and confirmed by western blotting to be at levels comparable to endogenous levels **(Fig. S1C)**. Twenty-four hours after replating, most of the junctions of the EGFP-afadin- expressing cells were linear and formed on the apical side. Immunofluorescence revealed that α-catenin and F-actin colocalized with apical linear junctions as in WT MDCK cells **(Fig. 2A)**. However, EGFP-afadinΔC did not localize apically but was diffusely distributed in the LCs. α-catenin and F-actin also diffusely localized on LCs, but did not accumulate in apical linear junctions, in a manner similar to afadinΔC. Furthermore, EGFP-afadinΔC-expressing cells were not rescued with respect to the degree of tilting and the lateral contact area of WT cells **(Fig. 1B&2A)**.

To further investigate the role of the C-terminal IDR of afadin in the localization of epithelial cells, we cultured the transfectants in collagen gel to allow for their growth into spherical cysts. Control EGFP-afadin-expressing cells formed typical epithelial cysts in which cells were arranged radially **(Fig. 2B)**. In the cysts, the localization of afadin is strictly localized in apical AJs; however, β-catenin and F-actin are found in both apical AJs and LCs, these differential localizations are similar to epithelial tissue than to 2D culture **(Fig. 2C)**. In contrast, although EGFP-afadinΔC-expressing cells formed single-lumen cysts, EGFP-afadinΔC failed to concentrate at the apical junction, the cells comprising the cysts were flattened, and these cell-cell junctions were tilted rather than radial **(Fig. 2B&S2A)**. The cell morphologies and structures of cell-cell junctions were remarkably similar to the results of the 2D culture. These results further confirm that the C-terminal IDR of afadin is important for apical localization and regulation of junction morphology.

Since nectins interact with afadin through their PDZ domain during AJ formation^24,25^, we examined whether forced expression of nectin alters afadin localization to the apical junctions. Nectin-1 and EGFP-afadin or EGFP-afadinΔC were stably expressed in afadin-KO cells **(Fig. 2D)**. Afadin and exogenous nectin-1 in EGFP- afadin expressing cells colocalized along apical junctions, hardly detected on LCs, with F-actin and α-catenin localized linearly along apical junctions **(Fig. 2D)**. In EGFP- afadinΔC-expressing cells, apical localization of afadin and nectin-1 was absent, and the linear signals of α-catenin and F-actin on apical junctions were not recovered.

Next, we expressed nectin-1 in afadin-KO cells and investigated the role of nectin-1 on AJ formation and the localization. To visualize nectin-1 in living cells, we stably expressed nectin-1 and the EGFP-tagged PDZ domain of afadin (AF_PDZ-EGFP) in WT or afadin-KO cells and observed the junction formation **(Fig. 2E&Videos S1, S2)**. The PDZ domain of afadin directly binds to the PDZ binding motif of nectins, reflecting the localization of nectins in the living cells. In afadin-KO cells, nectin-1 accumulated between cells, but its adhesion was unstable, and no apical localization was observed. These findings indicate that nectin binding to afadin alone is insufficient for apical junctional localization of afadin, and that nectin alone cannot accumulate in apical junctions.

### Destabilization of apical accumulation by lack of C-terminal IDR of afadin

To further analyze the role of the C-terminal disordered region of afadin during initial junction formation, we conducted live imaging of cells expressing EGFP-afadin and EGFP-afadinΔC **(Fig. 3A&Videos S3, S4)**. In the subconfluent cell layers, adherent cells extend and increase LC-like cell-cell contact. Live imaging showed that EGFP-afadin-expressing cells displayed convergence of cell-cell contacts to apical linear junctions, with EGFP-afadin concentrated linearly at the apical edge. In contrast, EGFP- afadinΔC signals were diffusely observed at LCs, displaying dynamic morphology changes but failing to form linear cell junctions within the observed timeframe **(Fig. 3A&Video S4)**. These results indicate that the C-terminal IDR of afadin is crucial for the convergence of cell-cell junctions from apical linear junctions. We also examined the effect of forced expression of nectin-1 in the accumulation of afadin in apical junction by live imaging and confirmed that EGFP-afadinΔC failed to accumulate at apical junctions even in the presence of nectin-1 **(Fig. 3B&Videos S5, S6)**.

**Figure 3.**
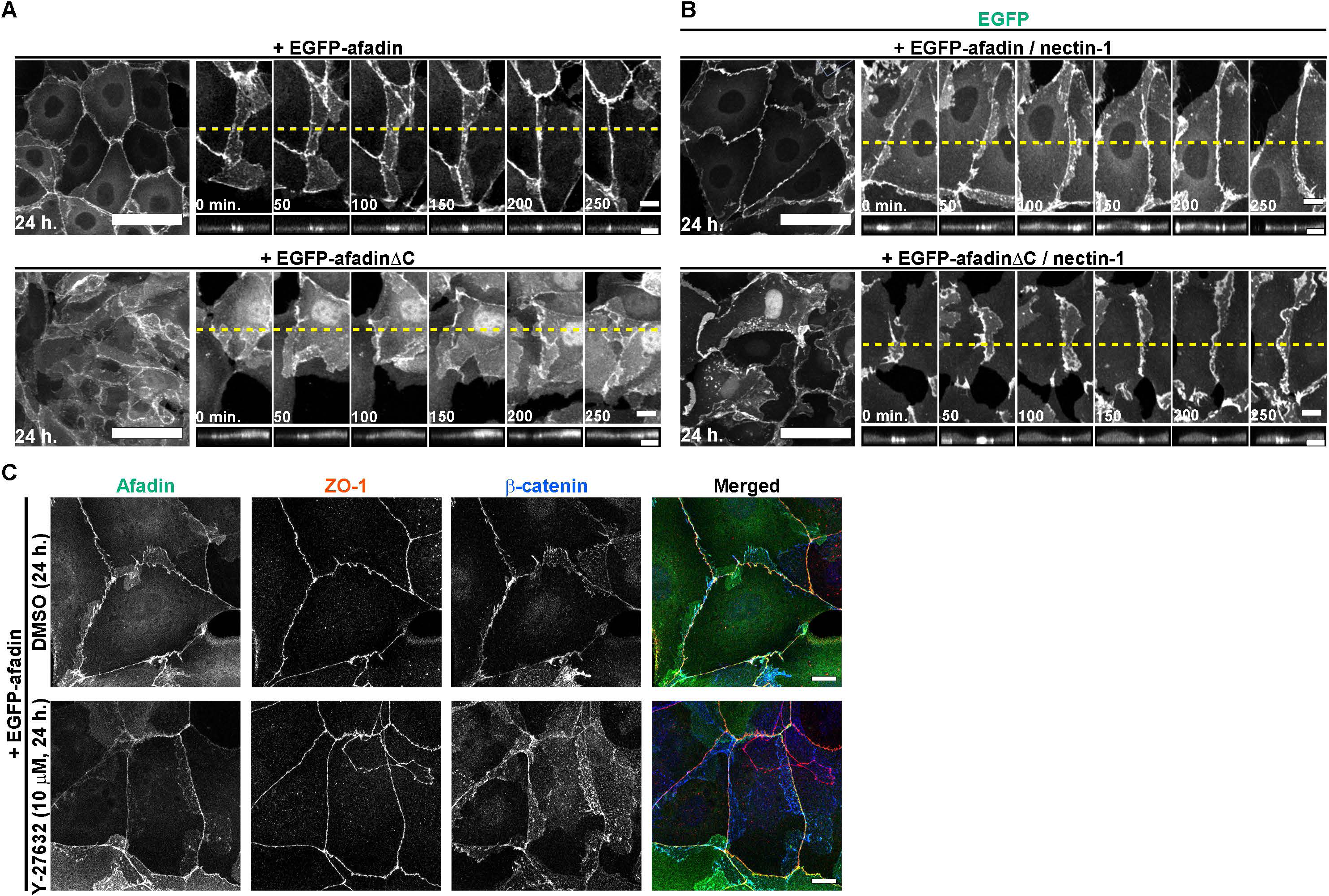
Localization of afadin to the apical junction is due to its self-organizing action **(A)** Junction formation of EGFP-afadin and EGFP-afadinΔC-expressing afadin-KO cells. (Left) Apical views of the transfectants cultured for 24 h. Scale bars, 50 μm. (Right) Time-lapse images of the junction formation; these cells were observed for 250 min at 50 min intervals. See also Videos S3&S4. Scale bars, 10 μm. (Bottom) Vertical sections of the upper images. Scale bars, 10 μm. **(B)** Junction formation of nectin-1 and EGFP-afadin- or EGFP-afadinΔC-expressing cells. The signals for EGFP-afadin or EGFP-afadinΔC are shown. (Left) Apical views of the transfectants cultured for 24 h. Scale bars, 50 μm. (Right) Time-lapse images of the junction formation; these cells were observed for 250 min at 50 min intervals. See also Videos S5&S6. Scale bars, 10 μm. (Bottom) Vertical sections of the upper images. Scale bars, 10 μm. **(C)** Distribution of afadin (green), ZO-1 (red), and β-catenin (blue) in EGFP-afadin-expressing cells upon the treatment of Y-27632. Apical views of the transfectants treated with DMSO or 10 μM Y-27632 for 24 h. A representative image is shown of six independent experiments. Scale bars, 10 μm.

Afadin regulates proper actomyosin association with AJs^15,26,27^. Therefore, we investigated whether myosin-induced contraction was involved in the apical accumulation of afadin during the initial junction formation **(Fig. 3C)**. When the cells were treated with Y-27632 from the beginning of culture^28^, afadin localized normally on the apical junctional complex within 24 h, indicating that myosin-dependent contractility was not involved in the apical accumulation of afadin during initial junction formation.

### No requirement of interaction with α-catenin or ZO-1 in apical localization of afadin

Previous studies have shown that the region deleted in afadinΔC contains the PR1-2 region and the CC domain necessary for binding to ZO-1 or α-catenin^29,30^. To confirm the binding of afadin to ZO-1 or α-catenin, we immunoprecipitated each EGFP-afadin fragment from the lysates of HEK293 transfectants and analyzed the co-precipitated proteins by western blotting, confirming that the PR1-2 region and CC domains of afadin were required for binding to ZO-1 and α-catenin, respectively **(Fig. S1E,F)**. Then, we examined the apical accumulation of EGFP-afadinΔC+CC in afadin-KO cells to investigate whether the CC domain of afadin is required for its apical junctional localization. EGFP-afadinΔC+CC partially localized to the apical junctions, and extensive lateral contact sites remained **(Fig. S2B)**. The F-actin organization of EGFP- afadinΔC+CC-expressing cells was similar to that of afadin-KO cells **(Fig. S2B)**. Live imaging showed that EGFP-afadinΔPR1-2, EGFP-afadinΔCC, but not EGFP- afadinΔC+CC accumulated in the apical junctions during initial junction formation **(Fig. S3A&Videos S7-9)**. These observations also suggest that the PR1-2 region and CC domain of afadin, which bind to ZO-1/2 and α-catenin, are not required for the apical localization of afadin.

To determine whether ZO-1/2 or αE-catenin is necessary for the apical localization of afadin, we examined its localization in ZO-1/2 double KO (dKO) cells and αE-catenin KO MDCK cells, which were previously generated^14^**(Fig. S1G,H)**. To assess the junctional localization of afadin, αE-catenin, and ZO-1 in ZO-1/2 dKO and αE-catenin KO cells, the cells were triple-stained with antibodies 24 h after replating **(Fig. S2D,E)**. In ZO-1/2 dKO cells, afadin was apically localized but showed a somewhat broader distribution in apical junctions, with a corresponding decrease in afadin localization **(Fig. S2D,F)**. In αE-catenin KO cells, despite unstable intercellular adhesions and reduced accumulation of afadin at cell-cell contacts, afadin and ZO-1 were colocalized at the apical side where cell-cell contacts formation was observed **(Fig. S2E, X-Z)**. These observations suggest that ZO-1/2 and α-catenin are involved in the apical localization of afadin; however, even in cells where either of them was KO, the apical localization of afadin was still maintained.

For a detailed analysis of the molecular behaviors of afadin and afadinΔC in ZO-1/2 dKO and αE-catenin-KO cells during initial junction formation, live imaging was conducted **(Fig. S3B,C&Videos S10-S13)**. EGFP-afadin-expressing ZO-1/2 dKO cells exhibited the convergence of spreading LCs to apical linear junctions, with EGFP-afadin concentrated linearly at the apical edge. In contrast, EGFP-afadinΔC-expressing ZO-1/2 dKO cells displayed diffuse signals at LCs, which failed to converge to linear cell junctions **(Fig. S3B&Video S10, S11)**. Live imaging of αE-catenin-KO cells revealed the accumulation of EGFP-afadin at the apical junctions. However, in EGFP-afadinΔC-expressing cells, partial linear localization of EGFP-afadinΔC signals and vigorously moving lateral contact sites were observed **(Fig. S3C&Videos S12, S13)**. These results suggest that ZO-1/2 or αE-catenin is not required for apical convergence of afadin from LCs.

### Implication of C-terminal IDR of afadin in condensates formation

Our observations suggest that the C-terminal region of afadin plays a role in the accumulation of apical junctions during initial junction formation, independent of its known binding partners. As mentioned previously, the C-terminal region of afadin is predicted to contain IDRs **(Fig. S4A)**. Increasing evidence suggests that multivalent protein-protein interactions mediated by IDRs facilitate the formation of condensates^30^. We transiently expressed EGFP-afadin in HEK293 cells to assess whether afadin forms IDR-dependent condensates. High EGFP-afadin expression resulted in the formation of large condensates **(Fig. 4A)**. We then transiently expressed various EGFP-afadin fragments in HEK293 cells to understand the role of each domain in condensate formation. EGFP-afadinΔRA, EGFP-afadinΔPDZ, EGFP-afadinΔPR1-2, EGFP- afadinΔCC, and EGFP-s-afadin all form condensates in the cells, similar to EGFP-afadin **(Fig. 4A)**. However, in EGFP-AF_N-PDZ- or EGFP-afadinΔC-expressing cells, the EGFP signal was higher in the cytoplasm, and condensate formation was significantly reduced compared to cells expressing EGFP-afadin **(Fig. 4A)**. Although higher expression induced the formation of large condensates, these observations suggested that afadin inherently possesses the capacity to form condensed compartments in an IDR-dependent manner in cells. Upon comparison of the results shown in Fig.4A and Fig. S1D, the localization of afadin to apical AJs was found to correspond to its ability to form condensates, suggesting that IDR-dependent condensate formation of afadin might be involved in the regulation of its localization. Subsequently, we constructed the EGFP- AF_1114-1654 fragment, containing only the IDR region deleted in afadinΔC **(Fig. S4A)**, and transiently expressed EGFP, EGFP-AF_1114-1654, and αE-catenin-EGFP in HEK293 cells **(Fig. 4B)**. EGFP-AF_1114-1654 expression resulted in the formation of abundant condensates; however, EGFP and αE-catenin-EGFP was not formed condensate **(Fig. 4B)**. We observed the dynamics of afadin condensates and found that these condensates were dynamic and fused within seconds in the cell, indicating liquid-like material properties (**Fig. 4C&Video S14**).

**Figure 4.**
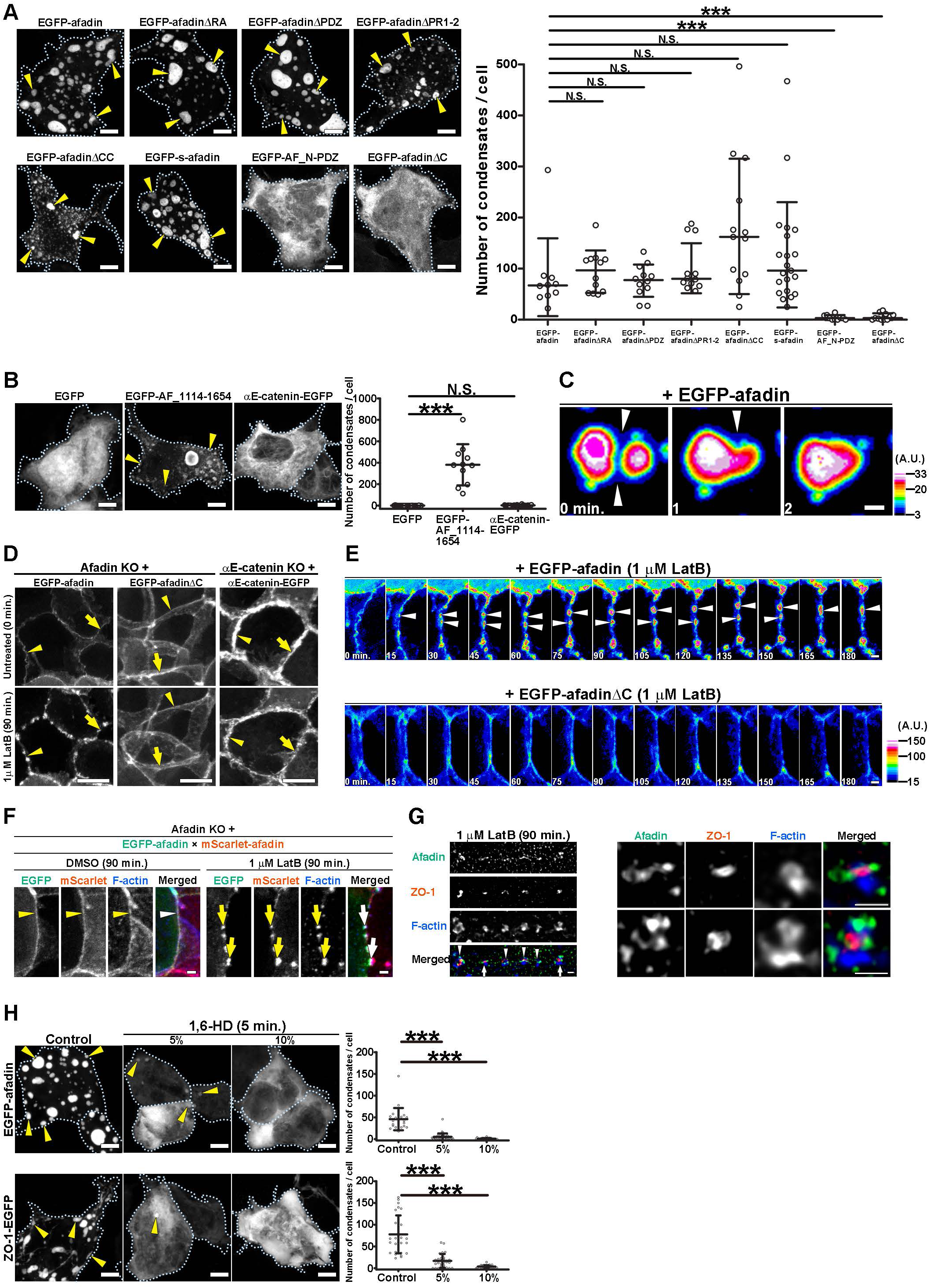
C-terminal region of afadin is implicated in condensates formation **(A)** Transient expression of EGFP-afadin, EGFP-afadinΔRA, EGFP-afadinΔPDZ, EGFP-afadinΔPR1-2, EGFP-afadinΔCC, EGFP-s-afadin, EGFP-AF_N-PDZ, and EGFP-afadinΔC in HEK293 cells; apical views of the transfectants cultured for 3 days. Dotted lines represent the cell of interest. Arrowheads indicate condensates. Scale bars, 5 μm. (Right) Quantification of EGFP condensate number in the cells. Whiskers represent the mean ± S.D. n = 10 for EGFP-afadin; n = 12 for EGFP-afadinΔRA; n = 12 for EGFP-afadinΔPDZ; n = 15 for EGFP-afadinΔPR1-2; n = 13 for EGFP-afadinΔCC; n = 21 for EGFP-s-afadin; n = 10 for EGFP-AF_N-PDZ; n = 11 for EGFP-afadinΔC. Steel’s multiple comparison test; N.S., not significant; ***, P ≤ 0.001. **(B)** Transient expression of EGFP, EGFP-AF_1114-1654, or αE-catenin-EGFP in HEK293 cells; apical views of the transfectants cultured for 3 days. Arrowheads indicate condensates. Scale bars, 5 μm. (Right) Quantification of EGFP condensate number of the cells. n = 48 for EGFP; n = 12 for EGFP-AF_1114-1654; n = 30 for αE-catenin-EGFP. Steel’s multiple comparison test; ***, P ≤ 0.001; N.S., not significant. **(C)** Fusion of afadin condensates in EGFP-afadin transiently expressing HEK293 cells. Apical views of condensates were observed for 2 min at 1 min intervals. Scale bar, 1 μm. The look-up table represents the intensity of EGFP signal. Arrowheads indicate point of fusion. A.U., arbitrary unit. See also Video S14. **(D)** Junction fragmentation in EGFP-afadin, EGFP- afadinΔC, and αE-catenin-EGFP expressing cells by treatment with LatB. (Upper) Apical views of the transfectants before treatment. (Bottom) Apical views of the transfectants after 90 min of treatment with 1 µM LatB. Arrowheads and arrows indicate the same junctions before and after treatment. Scale bars, 20 μm. **(E)** Live-cell imaging of junction fragmentation in EGFP-afadin or EGFP-afadinΔC expressing cells by the treatment of LatB. These cells were observed for 180 min at 15 min interval in the presence of 1 μM LatB. The look-up table represents the intensity of the EGFP signal. Scale bars, 2 μm. Arrowheads indicate condensates. See also Videos S15&S16. **(F)** Distribution of afadins in EGFP-afadin- or mScarlet-afadin-expressing afadin-KO MDCK cells upon treatment with LatB. Two transfectants were co-cultured for 3 days and treated with 1 μM LatB for 90 min. The cells were counterstained with phalloidin. Arrowheads indicate the apical cell-cell junctions. Arrows indicate fused condensates between adjacent cells. Scale bars, 2 μm. **(G)** (Left) Distribution of afadin (green), ZO- 1 (red) and F-actin (blue) in confluent monolayers of WT MDCK cells upon treatment with LatB and stained for the indicated molecules; apical views of the transfectants treated with DMSO or 1 μM LatB for 90 min. Arrows and arrowheads indicate condensates. The condensates indicated by arrows are shown in magnified images on the right. (Right) Projected magnified images of afadin, ZO-1, and F-actin in the condensates. A representative image is shown of four independent experiments. Scale bars, 1 μm. **(H)** Transient expression and distribution of EGFP-afadin or ZO-1-EGFP in HEK293 cells in the presence of 1,6-HD; apical views of the transfectants cultured for 3 days. Cells were treated with 1,6-HD for 5 min at 37°C. Arrowheads indicate condensates. Scale bars, 5 μm. (Right) Quantification of condensate number in the cells. Whiskers represent the mean ± S.D. n = 24 for EGFP-afadin without 1,6-HD; n = 43 for EGFP-afadin with 5% 1.6-HD; n = 23 for EGFP-afadin with 10% 1.6-HD; n = 30 for ZO-1-EGFP without 1,6-HD; n = 30 for ZO-1-EGFP with 5% 1,6-HD; n = 29 for ZO- 1-EGFP with 10% 1,6-HD. Steel’s multiple comparison test; ***, P ≤ 0.001.

To examine the impact of F-actin polymerization disassembly on the molecular behavior of afadin already localized at cell-cell junctions, we treated subconfluent afadin-KO MDCK cells expressing EGFP-afadin or EGFP-afadinΔC with latrunculin-B (LatB), which reversibly blocks actin polymerization. EGFP-afadin was initially linearly localized at the apical junctions; however, after 90 min of LatB treatment, continuous junctional afadin was segmented into a series of condensates **(Fig. 4D)**. However, the localization of afadinΔC was unaffected by LatB treatment, and no condensates were observed **(Fig. 4D)**. As a control, we examined the localization of αE-catenin-EGFP in αE-catenin-KO cells following LatB treatment. In αE-catenin-EGFP-expressing cells, some junctional αE-catenin-EGFP was partially fragmented from its linear localization 90 min following LatB treatment, but no condensates were observed. We also investigated the amount of residual F-actin present at the junctions in our experiments and confirmed the effectiveness of LatA **(Fig. S4B)**. Live-cell imaging revealed that the formation of afadin condensates followed the fragmentation of junctional afadin, which actively moved and fused into condensates **(Fig. 4E&Video S15)**. However, in EGFP- afadinΔC-expressing cells, these behaviors were not observed, and junctional afadin remained stable following LatB treatment **(Fig. 4E&Video S16)**. Then, we examined the effect of F-actin inhibition on afadin condensate formation using HEK293 cells **(Fig. S4C)**. We found that condensates were still formed; however, analysis of condensate size and shape in LatB-treated cells revealed that these condensates took more irregular shapes compared to DMSO-treated control cells. In addition, s-afadin, which lacks the F-actin binding region, forms condensates as well as full-length afadin **(Fig. 4A)**. These results indicate that the binding of afadin to actin alters the morphology of the condensates but was not essential for condensate formation.

Next, to investigate the formation of condensates from junctional afadin, we examined the molecular behavior in a mixed culture of EGFP-afadin- and mScarlet- afadin-expressing afadin-KO MDCK cells **(Fig. 4F)**. In the heterotypic cell-cell junctions between EGFP-afadin- and mScarlet-afadin-expressing cells, EGFP-afadin, mScarlet-afadin, and F-actin colocalized, suggesting the normal formation of AJs between these cells. In contrast, in the presence of LatB for 90 min, these signals were observed in the condensates at cell junctions. Interestingly, the F-actin signal was observed throughout the individual condensates, whereas the EGFP and mScarlet signals were symmetrical on both sides of the condensates. This result indicates that afadin, which forms condensates in individual cells, behaves as a dynamic condensate when linked between cells.

To understand the relationship between the molecular interactions and condensate formation of endogenous afadin and ZO-1, we examined the localization of afadin, ZO-1, and F-actin in WT MDCK cells in the presence of LatB using immunofluorescence staining **(Fig. 4G)**. After 90 min of LatB treatment, endogenous afadin, ZO-1, and F-actin were fragmented at the cell junctions and had a condensate-like morphology, consistent with the observations of afadin in Fig. 4D. To examine the localization of afadin, ZO-1, and F-actin in more detail, cell junctions were observed at a high magnification using super-resolution microscopy **(Fig. 4G, Right)**. The condensate-like structures of endogenous afadin and ZO-1 were adjacent to each other without colocalization, suggesting that these condensate-like structures are segregated in the juxta-membrane region. On the other hand, instances of F-actin co-localization with afadin and ZO-1 were also observed; however, these three molecules appeared to remain segregated within the aggregate. This suggests that the condensates of ZO-1, which are thought to act as the constitutive hubs of TJs, and those of afadin, function in different fractions. In summary, the results suggest that afadin condensates not only exhibit liquid-like properties in AJs but also function intercellularly in a manner that is distinct from the TJ phase-separated compositional function of ZO-1.

We then transiently expressed EGFP-afadin in ZO-1/2 dKO MDCK cells to determine whether ZO proteins were required for afadin to form condensates^16^. Previous studies have shown that exogenous afadin is enriched in the condensed ZO-1 phase of HEK293 cells. We observed the formation of EGFP-afadin condensates in ZO-1/2 dKO cells **(Fig. S4D)**. These results indicated that EGFP-afadin condensates were formed in a ZO-independent manner.

To investigate the properties of afadin condensate formation in HEK293 cells, we treated the cells with 1,6-hexanediol (HD). The aliphatic alcohol 1,6-HD has been widely used to study the formation process of membrane-free condensates by liquid- liquid phase separation (LLPS). 1,6-HD inhibits the weak hydrophobic protein-protein interactions required for the formation of liquid-like condensates^31,32^. Since ZO-1 has been reported to form condensates induced by overexpression^16–18^, we used ZO-1-EGFP as a control. Treatment with 1,6-HD of cells expressing EGFP, EGFP-afadin, and ZO-1- EGFP showed that 5% and 10% 1,6-HD reduced the condensates of afadin and ZO-1 within 5 minutes in a concentration-dependent manner, suggesting that these condensates might be formed through LLPS driven by weak hydrophobic protein– protein interactions **(Fig. 4H)**.

### Liquid-like properties of afadin promote condensate formation and junctional accumulation

Building on the results of Fig. 4, which suggest that afadin forms condensates via LLPS, we employed fluorescence recovery after photobleaching (FRAP) and fluorescence correlation spectroscopy (FCS) to analyze the molecular dynamics of afadin at the apical junction and within the cytoplasm **(Fig. 5A-C)**. First, we performed FRAP analysis on the condensates in HEK293 cells, and partially bleached EGFP-afadin condensates and measured recovery times **(Fig. 5A&Video S17)**. Afadin condensates showed rapid recovery within seconds, indicating liquid-like material properties. Next, we analyzed the dynamics of mScarlet-afadin and mScarlet-afadinΔC at the apical junction in MDCK cells. We found that mScarlet-afadin recovered up to 50% of its initial concentration at 37°C with a time constant of t1/2 = 25.3 ± 4.9 s and that mScarlet-afadinΔC recovery was faster with t1/2 = 19.8 ± 5.5 s **(Fig. 5B&Videos S18, S19)**.

**Figure 5.**
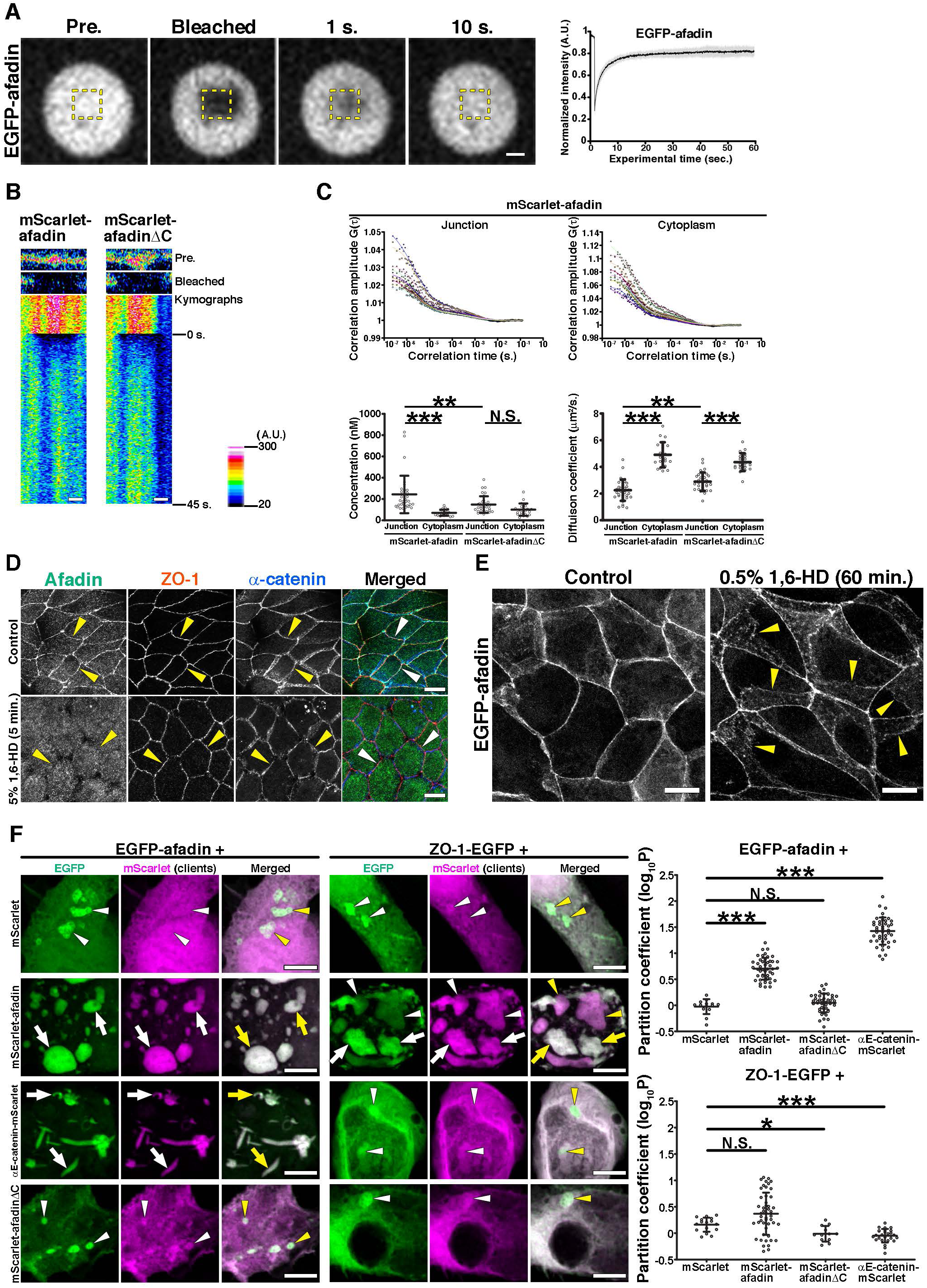
Liquid-like properties of afadin promote junctional accumulation **(A)** Fluorescence recovery after photobleaching (FRAP) measurements of EGFP-afadin in condensates within HEK293 cells. Dashed boxes indicate the bleached region. Scale bar, 1 μm. See also Video S17. (Right) Quantification of the region of interest of EGFP signal recovery. A line represents the mean ± S.D. A.U., arbitrary unit. n = 5. **(B)** mScarlet-afadin and mScarlet-afadinΔC dynamics at the junctions during FRAP. mScarlet-afadin or mScarlet-afadinΔC expressing ZO-1/2 dKO MDCK cells were selectively bleached at the apical junctions, and recovery was measured over time at 37°C. See also Videos S18&S19. Kymographs show the fluorescence intensities of both mScarlet-afadin and mScarlet-afadinΔC at the apical junctions. The look-up table represents the intensity of the mScarlet signal. Scale bars, 2 μm. **(C)** Fluorescence correlation spectroscopy (FCS) analysis of the molecular dynamics of afadin. (Upper) FCS analysis of mScarlet-afadin at apical junctions and in the cytoplasm. Correlation curves were fitted using a two-component 3D diffusion model. Data were collected from 9 different cells for mScarlet-afadin at apical junctions (left) and in the cytoplasm (right). (Bottom) Summary of FCS quantification. Molecular concentrations (left) and diffusion coefficients (right) of mScarlet-afadin and mScarlet-afadinΔC at apical junctions and in the cytoplasm. n = 33 for mScarlet-afadin and mScarlet-afadinΔC at junctions; n = 22 for mScarlet-afadin and mScarlet-afadinΔC in the cytoplasm. Steel and Dwass’s multiple comparison test; ***, P ≤ 0.001; **, P ≤ 0.01; N.S., not significant. **(D)** Distribution of afadin (green), ZO-1 (red), and α-catenin (blue) of WT MDCK cells upon treatment with 1,6-HD; apical views of the transfectants treated with or without 5% 1,6- HD for 5 min at 37°C and stained for the indicated molecules. Arrowheads indicate the apical cell junctions. Scale bars, 10 μm. **(E)** Distribution of EGFP-afadin of afadin-KO cells upon treatment with 1,6-HD; apical views of the transfectants treated with or without 0.5% 1,6-HD for 60 min at 37°C. Arrowheads indicate the apical cell junctions. Scale bars, 10 μm. **(F)** (Left) Partitioning coefficient assay of afadin and ZO-1. Transient co-expression of EGFP-afadin or ZO-1-EGFP (green) and mScarlet fusion clients (magenta) in HEK293 cells; apical views of the transfectants cultured for 3 days. Arrowheads indicate droplets with EGFP signal. Arrows indicate colocalized droplets with EGFP and mScarlet signals. A representative image is shown of six independent experiments. Scale bars, 5 μm. (Right) Quantification of partitioning coefficient (P) of mScarlet signals in each droplet. Graphs were visualized as log10P. A log10P value of 0 reflects equal partitioning of condensate and cytoplasm, while values higher than 0 indicate more clients in the condensate and lower than 0 indicate more in the cytoplasm. Whiskers represent the mean ± S.D. n = 14 for EGFP-afadin/mScarlet. n = 46 for EGFP-afadin/mScarlet-afadin. n = 43 for EGFP-afadin/αE-catenin-mScarlet. n = 44 for EGFP-afadin/mScarlet-afadinΔC. n = 14 for ZO-1-EGFP/mScarlet. n = 46 for ZO-1- EGFP/mScarlet-afadin. n = 43 for ZO-1-EGFP/αE-catenin-mScarlet. n = 44 for ZO-1- EGFP/mScarlet-afadinΔC. Steel’s multiple comparison test; ***, P ≤ 0.001; *, P ≤ 0.05; N.S., not significant.

We then used FCS to measure the concentration and diffusion coefficient of mScarlet-afadin and mScarlet-afadinΔC. FCS analysis at the apical junction and in cytoplasm identified that these two molecules have different concentrations and diffusion coefficients in the AJ and in the cytoplasm, respectively **(Fig. 5C)**. Molecular concentrations of mScarlet-afadin were significantly higher in AJ than in cytoplasm. Furthermore, the diffusion coefficient of mScarlet-afadin differed between the AJ and cytoplasm. The diffusion coefficient of afadin in the AJ was reduced to about half of that in the cytoplasm. In contrast, afadinΔC showed a much smaller decrease in its diffusion coefficient between the AJ and the cytoplasm **(Fig. 5C)**. Molecular diffusion within the condensed domain is shown to be reduced when LLPS is occurring^33,34^, and these observations support the idea that afadin is in a different state due to the LLPS between the AJ and cytoplasm.

To investigate whether LLPS plays a role in the localization of afadin to apical AJs, we treated WT MDCK cells with 1,6-HD for 5 min and subsequently examined the localization of afadin, ZO-1, and α-catenin **(Fig. 5D)**. We found that endogenous afadin localization disappeared from AJs, while ZO-1 and α-catenin still localized in the junctions in a linear fashion **(Fig. 5D)**. This result indicates that afadin lost its localization to AJs in a 1,6-HD-dependent manner. In contrast, ZO-1 and α-catenin did not lose their localization, suggesting that the apical localization of afadin is not dependent on its binding to ZO-1 or α-catenin. Furthermore, when EGFP-afadin- expressing cells forming apical AJs were treated with low concentrations of 1,6-HD for 60 min, the localization of apical AJs was reduced and adhesive structures resembling of the afadinΔC-expressing cells were observed **(Fig. 5E)**. These observations suggest that afadin forms condensates in the nature of LLPS and that this property is responsible for its accumulation in apical AJs.

As shown in Fig. 4G, endogenous afadin and ZO-1 formed distinct condensates. To investigate this further, we conducted co-expression experiments with different pairs of afadin, ZO-1, and αE-catenin in HEK293 cells **(Fig. 5F)**. When EGFP-afadin and mScarlet-afadin were co-expressed, both proteins were evenly distributed. However, when afadin and ZO-1 were co-expressed, the two proteins formed droplets but were not evenly distributed **(Fig. 5F, Right)**. Specifically, the distribution coefficients showed a large standard deviation, indicating that afadin and ZO-1 were not uniformly distributed, suggesting partial segregation within the same condensate. This inhomogeneity contrasts with the control experiment of EGFP-afadin and mScarlet-afadin, supporting the idea that afadin and ZO-1 differ in their spatial organization within the condensate. These observations suggest that afadin and ZO-1, both of which exhibit LLPS properties, are distributed differently, even within the same condensate.

On the other hand, when afadin and αE-catenin were co-expressed, afadin co- localized with αE-catenin in the cytoplasm and formed fibrous, irregularly shaped condensates. Since αE-catenin does not exhibit LLPS-mediated condensate formation **(Fig. 4B)** and the morphology of this condensate is not droplet-like, it is considered an aggregate formed by the binding of afadin and αE-catenin. In contrast, when ZO-1 and αE-catenin were co-expressed, both molecules were evenly distributed in the cytoplasm, but only ZO-1 formed condensates, which lacked αE-catenin. We also performed co- expression experiments with afadinΔC and found that afadinΔC did not affect the condensate formation of the partner molecule. These results suggest that heterogeneity in their co-condensation, potentially influenced by the presence or absence of LLPS property or in the interacting proteins.

### Multivalent afadin interaction promotes IDR-mediated junction accumulation

So far, it has been found that the C-terminal IDR of afadin is essential for condensate formation through LLPS and for its localization to the apical AJ. However, fragments containing only the IDR fail to localize to the apical AJ **(Fig. S4E)**, suggesting that domains other than the IDR are also required. To investigate this, we generated various afadin fragments that include the RA, DIL, PDZ, and FAB domains, along with the IDR, and assessed their localization **(Fig. S5A)**. These fragments were expressed in afadin- KO MDCK cells, and their localization was subsequently analyzed **(Fig. 6A)**. Among these molecules, those containing both FAB and IDR, along with either the RA or PDZ domains, were localized apically, whereas molecules lacking the FAB domain showed reduced localization at the AJ. These results indicate that the apical localization of afadin requires multivalent interactions with cell adhesion molecules and the cytoskeleton, including IDRs. Next, we generated various chimeric molecules by combining the RA, DIL, PDZ, and FAB domains of afadin with IDRs derived from ZO-1 and evaluated their localization **(Fig. 6B&Fig. S5B)**. Notably, the C-terminal IDR of ZO-1 (ZO-1_C) does not localize to cell junctions as previously reported^35^. However, our chimeric molecules showed apical localization, indicating that differences in IDRs play a critical role in the localization properties of these molecules.

**Figure 6.**
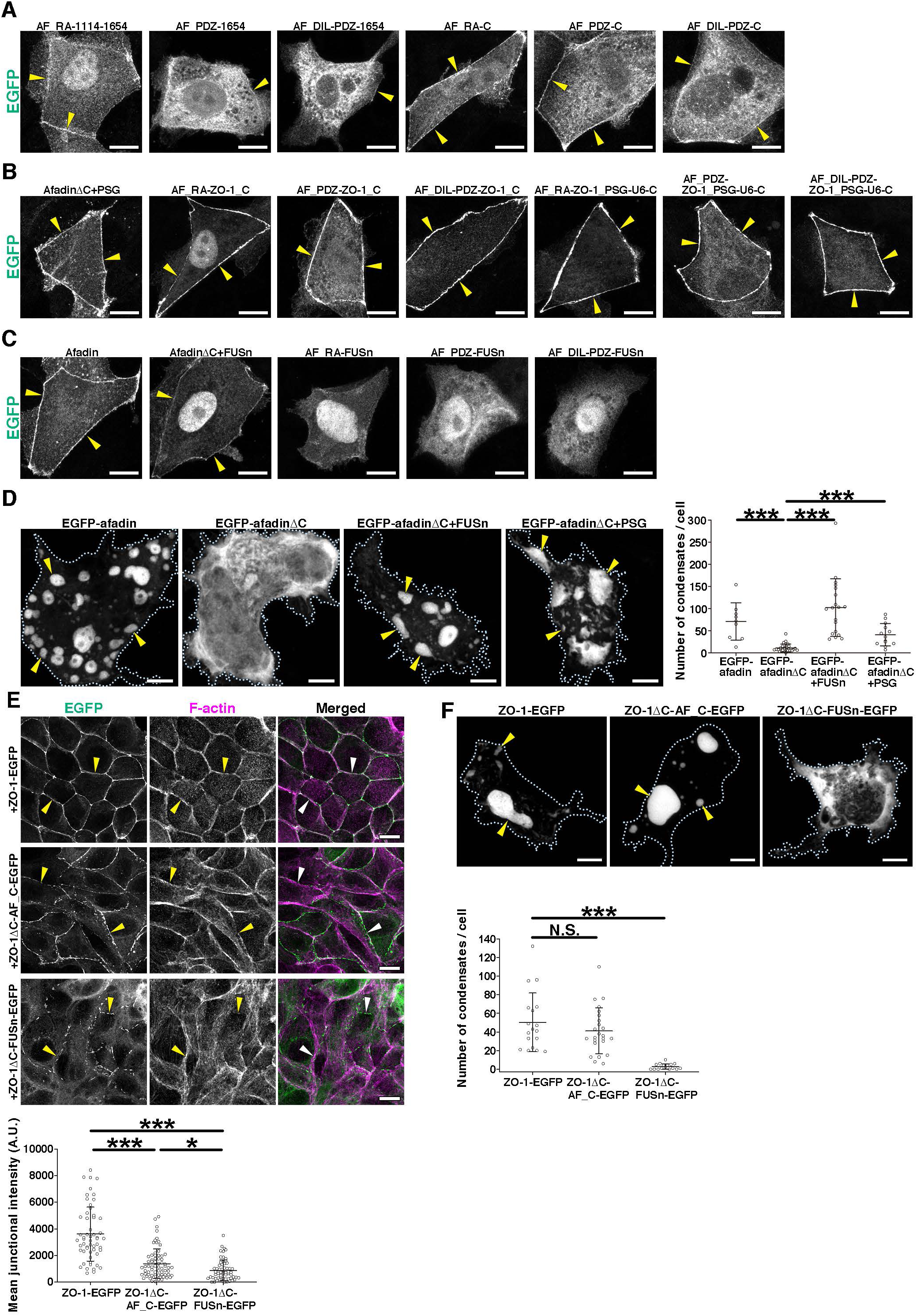
Different IDRs of afadin alter localization within AJC **(A)** Distribution of various afadin fragments. Scale bars, 10 μm. Arrowheads indicate the junctional EGFP signals. See also Fig. S5A. **(B)** Distribution of afadin-ZO-1 chimeras. Scale bars, 10 μm. See also Fig. S5B. **(C)** Distribution of afadin-FUSn chimeras. Scale bars, 10 μm. See also Fig. S5C. **(D)** Transient expression of EGFP- afadin, EGFP-afadinΔC, EGFP-afadinΔC+FUSn, or EGFP-afadinΔC+PSG in HEK293 cells; apical views of the transfectants cultured for 3 days. Dotted lines represent the cell of interest. Arrowheads indicate condensates. Scale bars, 5 μm. (Right) Quantification of EGFP condensate number in the cells. Whiskers represent the mean ± S.D. n = 9 for EGFP-afadin; n = 28 for EGFP-afadinΔC; n = 19 for EGFP-afadinΔC+FUSn; n = 13 for EGFP-afadinΔC+PSG. Steel and Dwass’s multiple comparison test; ***, P ≤ 0.001. **(E)** (Upper) Distribution of ZO-1 (green) and F-actin (magenta) in ZO-1-EGFP, ZO-1ΔC- AF_C-EGFP, or ZO-1ΔC-FUSn-EGFP-expressing ZO-1/2 dKO cells; apical views of the transfectants cultured for 24 h and stained for the indicated molecules. Arrowheads indicate apical junctions. Scale bars, 10 μm. (Bottom) Quantification of junctional EGFP signal. Junctional signal acquired at 5 μm^2^. Cytoplasmic signals averaged 5 areas of 25 μm^2^. Junctional signal subtracted cytoplasmic signal. A.U., arbitrary unit. Whiskers represent mean ± S.D. n = 55 for ZO-1-EGFP. n = 60 for ZO-1ΔC-AF_C-EGFP. n = 61 for ZO-1ΔC-FUSn-EGFP. Steel and Dwass’s multiple comparison test; ***, P ≤ 0.001. *, P ≤ 0.05. **(F)** Transient expression of ZO-1-EGFP, ZO-1ΔC-AF_C-EGFP, and ZO- 1ΔC-FUSn-EGFP in HEK293 cells; apical views of the transfectants cultured for 3 days. Arrowheads indicate condensates. Scale bars, 5 μm. (Bottom) Quantification of EGFP condensate number in the cells. Whiskers represent the mean ± S.D. n = 18 for ZO-1-EGFP. n = 24 for ZO-1ΔC-AF_C-EGFP. n = 16 for ZO-1ΔC-FUSn-EGFP. Steel’s multiple comparison test; ***, P ≤ 0.001; N.S., not significant.

### Apical accumulation of chimeric afadin of FUS-derived IDR and afadinΔC

Our results indicate that the nature of IDRs causes the apical accumulation of afadin. If the localization of afadin is regulated by IDRs, it is conceivable that the IDRs of other molecules may also be responsible for its apical localization. Although several proteins are involved in IDR-dependent condensate formation, the RNA-binding protein Fused in Sarcoma (FUS) is one of the most studied proteins in terms of its LLPS properties^36–38^. The low-complexity N-terminal IDR of FUS (FUSn) is responsible for its structural disorder in solution and formation of a liquid-like phase-separated state. To test this idea, we constructed EGFP-afadinΔC+FUSn, which consists of an FUSn sequence added to the C-terminal side of the EGFP-afadinΔC fragment **(Fig. S5C, See also STAR methods)**. We then stably expressed EGFP-afadinΔC+FUSn fragments in afadin-KO cells to examine the effect of the FUSn sequence on afadinΔC localization. EGFP- afadinΔC+FUSn localized linearly along the apical junctions, and the localization of LCs was drastically reduced **(Fig. 6C)**. The IDR sequences in FUS have been shown to be involved in condensate formation^39^. We subsequently created chimeric molecules in which each domain of afadin was bound to FUSn sequence and examined whether these molecules localized to apical AJ **(Fig. 6C)**. Chimeric molecules in which afadin’s IDR was replaced with a FUS-derived IDR localized apically, whereas chimeric molecules containing only a single domain of afadin combined with a FUS-derived IDR failed to localize apical AJ. To confirm the properties of EGFP-afadinΔC+FUSn, we transiently expressed this fragment in HEK293 cells and observed condensate formation **(Fig. 6D)**. This indicated that FUSn was responsible for condensate formation. The FUSn sequence is not known to interact with any cell adhesion-related molecule, and the addition of this sequence resulted in the apical localization of afadin, suggesting that IDRs are required for the accumulation of afadin at the apical junctions of cells.

We further examined the effect on localization of replacing the IDR of ZO-1 with either an afadin- or FUS-derived IDR **(Fig. 6E&Fig. S5D)**. These chimeric molecules were often localized at the apical junctions, but their localization was not as fully restored as that of the full-length molecules, and a decrease in continuous localization at the junctions was observed. The localization of these chimeric molecules to apical AJs corresponded to some extent with their ability to form droplets, and ZO- 1ΔC_FUSn-EGFP that were unable to form droplets showed minimal localization to AJs **(Fig. 6F)**.

### Altered distribution of afadin within the AJC by different IDR

As summarized in **Fig. 7A**, replacing the IDR of afadin with the IDR of another molecule restored localization to apical AJ. To obtain more detailed information about the distribution of EGFP-afadinΔC+FUSn and EGFP-afadinΔC+PSG, we immunostained cells expressing EGFP-afadin, EGFP-afadinΔC+FUSn, and EGFP-afadinΔC+PSG with ZO-1 and β-catenin and observed the cells using super-resolution confocal microscopy. EGFP-afadin colocalized with β-catenin and ZO-1 at the AJCs, and these molecules appeared to be closely associated without apparent separation. Although EGFP- afadinΔC+FUSn and EGFP-afadinΔC+PSG were also localized to the AJCs, its distribution differed from that of EGFP-afadin; the localization of these chimeric molecules was shifted away from the ZO-1 and β-catenin in both X-Y and X-Z planes **(Fig. 7C)**. The distribution of these molecules was assessed by line scan, which revealed a minimal shift in localization between ZO-1 and β-catenin for EGFP-afadin, contrasting with a substantial shift observed for EGFP-afadinΔC+FUSn **(Fig. 7C)**. Furthermore, a reduced frequency of colocalization was observed with ZO-1 and β-catenin, and the colocalized regions of these molecules were significantly reduced **(Fig. 7C)**. These observations indicate that the FUS-derived IDR added to afadinΔC allows it to localize apically **(Fig. 7B, X-Z and Y-Z)**. However, the distinct distribution of EGFP- afadinΔC+FUSn and EGFP-afadinΔC+PSG compared to EGFP-afadin suggests that the different IDRs may have resulted in a different distribution within the AJC.

**Figure 7.**
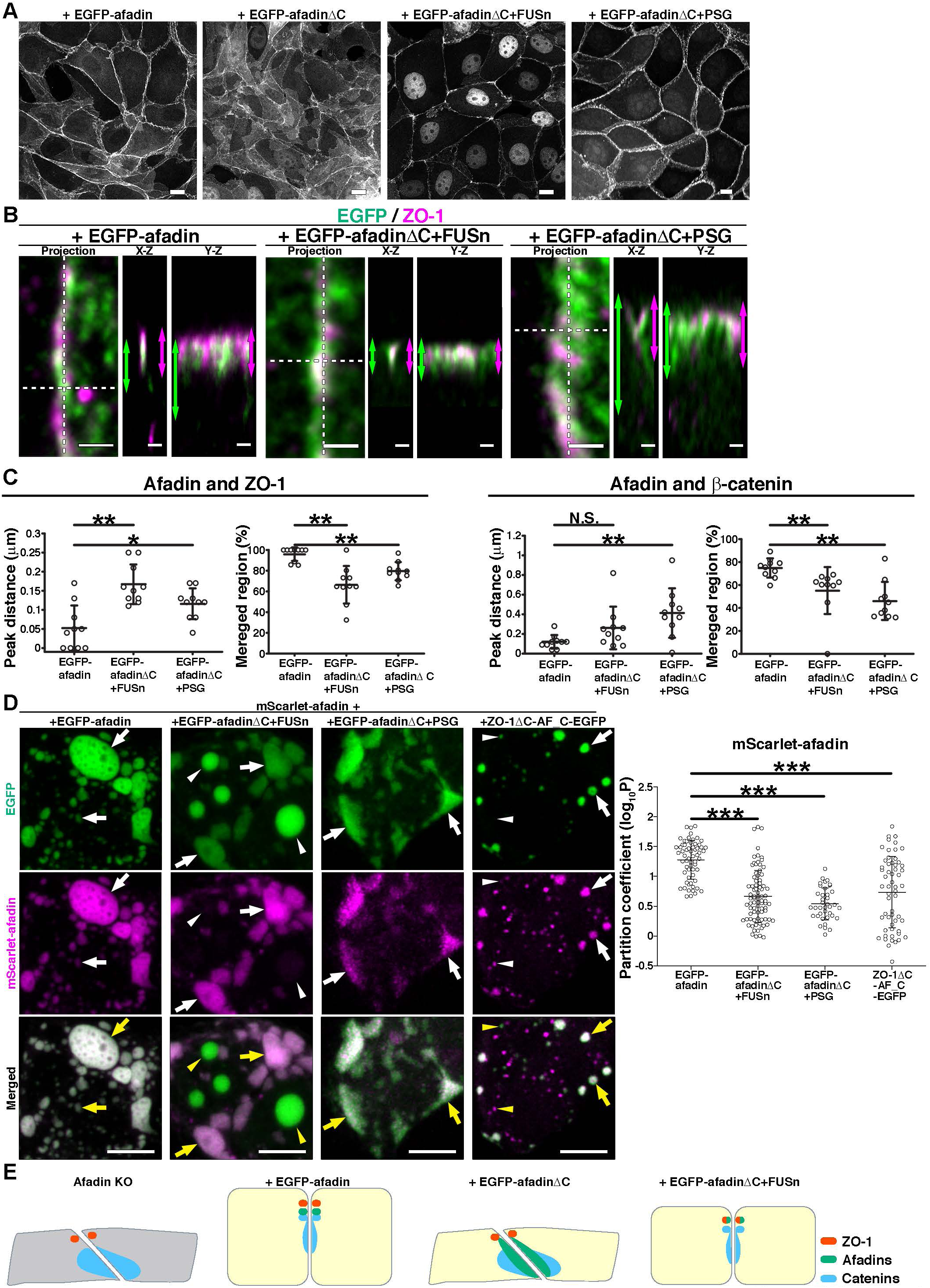
Different IDRs of afadin alter localization within AJC **(A)** Localizations of EGFP-afadin-, EGFP-afadinΔC-, EGFP-afadinΔC+FUSn-, or EGFP-afadinΔC+PSG expressing afadin-KO cells; apical views of the transfectants cultured for 24 h. Scale bars, 10 μm. **(B)** Close up views of the junctional distribution of afadins (green) and ZO-1 (magenta) in EGFP-afadin-, EGFP-afadinΔC+FUSn-, or EGFP-afadinΔC+PSG-expressing afadin-KO cells; apical views of the transfectants cultured for 24 h and visualized with the indicated molecules. Dashed lines indicate the position of vertical section. (X-Z) Vertical sections of the projection images. Bidirectional arrows are the position of the signal detection point. (Y-Z) Vertical sections of the projection images. Scale bars, 0.5 μm. **(C)** Quantification of peak distance and merged region between afadins and ZO-1 or β-catenin in EGFP-afadin-, EGFP- afadinΔC+FUSn-, or EGFP-afadinΔC+PSG-expressing afadin-KO cells. Whiskers represent the mean ± S.D. Peak distance between afadins and ZO-1; n = 10 for EGFP- afadin, n = 10 for EGFP-afadinΔC+FUSn, n = 10 for EGFP-afadinΔC+PSG. Merged region with afadins and ZO-1; n = 10 for EGFP-afadin, n = 10 for EGFP- afadinΔC+FUSn, n = 10 for EGFP-afadinΔC+PSG. Peak distance between afadins and β-catenin; n = 10 for EGFP -afadin, n = 10 for EGFP-afadinΔC+FUSn, n = 10 for EGFP- afadinΔC+PSG. Merged region with afadins and β-catenin; n = 10 for EGFP-afadin, n = 10 for EGFP-afadinΔC+FUSn, n = 10 for EGFP-afadinΔC+PSG. Steel’s multiple comparison test; **, P ≤ 0.01; *, P ≤ 0.05; N.S., not significant. **(D)** Partitioning coefficient assay of afadin and ZO-1 chimeras. Transient co-expression of EGFP tagged afadin or chimeras (green) and mScarlet-afadin clients (magenta) in HEK293 cells; apical views of the transfectants cultured for 3 days. Arrowheads indicate condensates consisting primarily of a single molecular species. Arrows indicate condensates consisting of mixture of two molecular species. Scale bars, 5 μm. (Right) Quantification of partitioning coefficient (P) of mScarlet signals in each droplet. Graphs were visualized as log10P. A log10P value of 0 reflects equal partitioning of condensate and cytoplasm, while values higher than 0 indicate more clients in the condensate and lower than 0 indicate more in the cytoplasm. Whiskers represent the mean ± S.D. n = 62 for EGFP- afadin/mScarlet-afadin. n = 81 for EGFP-afadinΔC+FUSn/mScarlet-afadin. n = 38 for EGFP-afadinΔC+PSG/mScarlet-afadin. n = 57 for ZO-1ΔC-AF_C-EGFP/mScarlet- afadin. Steel’s multiple comparison test; ***, P ≤ 0.001. **(E)** Schematic illustrations of junctional structures of afadin-KO cells and EGFP-afadin-, EGFP-afadinΔC-, and EGFP-afadinΔC+FUSn-expressing afadin-KO cells.

As shown in Figs.4G&5F, afadin and ZO-1 were differentially distributed in the condensates, suggesting that these condensates were not mixed. Furthermore, because differences in the IDR regions cause differences in the localization of afadin in Fig.7B, we considered the possibility that differences in IDRs regulate molecular localization within the AJC. To test this hypothesis, we constructed EGFP-afadinΔC+PSG, which consists of the PDZ3-SH3-GuK (PSG) sequence of ZO-1 on the C-terminal side of the EGFP-afadinΔC fragment **(Fig. S5B, See also STAR methods)** because the PSG domain of ZO-1 is known to localize to the initial AJ through dimerization between ZO-1^40^. To confirm the properties of EGFP-afadinΔC+PSG, we transiently expressed this fragment in HEK293cells and observed condensate formation **(Fig. 6F)**. We then stably expressed EGFP-afadinΔC+PSG in afadin-KO cells to examine their localization in the AJC. The localization of EGFP-afadinΔC+PSG was similar to that of EGFP- afadinΔC+FUSn, with linear localization to AJC, while there was little localization to the lateral contacts **(Fig. 7B)**. The distribution of EGFP-afadinΔC+PSG was quantified by a line scan, which revealed a significant peak shift in localization between ZO-1 and β-catenin compared to EGFP-afadin **(Fig. 7C)**. However, the localization of EGFP- afadinΔC+PSG was close to ZO-1 and far from β-catenin. Interestingly, EGFP- afadinΔC+PSG and ZO-1 were separated from each other and localized in a double- linear manner within the apical junction complex **(Fig. 7B)**.

To investigate this further, we conducted co-expression experiments of mScarlet-afadin and EGFP-afadinΔC+FUSn or EGFP-afadinΔC+PSG in HEK293 cells **(Fig. 7D)**. When mScarlet-afadin and EGFP-afadinΔC+FUSn were co-expressed, the two proteins formed droplets but were not uniformly distributed, rather they formed different droplets **(Fig. 7D)**. These observations suggest that differences in the nature of IDRs regulate their distribution within the AJC in epithelial cells **(Fig. 7E)**.

## Discussion

Correct positioning of the AJ is considered a crucial step in establishing apical-basal polarity and TJ formation in epithelial cells. Afadin and nectins are strictly localized to AJs in epithelial cells; however, the mechanism underlying this localization remains unknown. Afadin-KO mice exhibit impaired AJ formation, disruption of apical-basal polarity, and TJ formation in the neuroepithelium of the ectoderm^10,11^. Furthermore, experiments using afadin-KO cells demonstrate that afadin is essential for cell intercalation during mosaic-like cell patterning **(Fig. 1E)**. The abnormalities observed in afadin-KO mice and cells are likely due to the failure of AJs to form at the correct locations. Despite its recognized importance, the precise mechanisms by which afadin regulates intercellular adhesion and polarity remain elusive. The present study revealed that afadin accumulates at the apical junction of cells, driven by the nature of its IDRs, independent of known domains during initial junction formation. Deletion of the afadin IDR suppressed its apical accumulation and concomitantly reduced the accumulation of the cadherin/catenin complex and F-actin. When the IDR fragment of afadin was overexpressed in cells, it formed condensates; however, IDR alone did not accumulate at the apical junction **(Fig. S4D)**. Intriguingly, replacing the IDR of afadin with the IDR of FUS restored apical localization but altered the distribution of afadin within the AJC. These results indicate that IDR is necessary not only for apical localization but also for regulating their distribution within the AJC. The IDR at the C-terminus of afadin alone is not sufficient for junctional localization, and the N-terminus of afadin without the IDR cannot successfully converge to the apical junctions. When afadin accumulates in apical AJs, multivalent interactions through the adhesion molecules and the cytoskeleton may trigger IDR-mediated condensate formation, which may facilitate efficient accumulation.

### IDR-dependent apical localization of afadin

Previously, Takai et al. reported that nectin and cadherin are diffusely distributed on the free surface of the membranes of migrating cells at the initial stage of AJ formation. When two migrating cells contact each other through protrusions such as filopodia and lamellipodia, nectin first forms trans-dimers at intercellular contact sites, and nectin/afadin complexes recruit cadherin to the initial contact sites. Subsequently, nectin/afadin gradually localizes towards the apical side, resulting in the linear localization of the actin cytoskeleton and cadherin/catenin complexes at the apical side, eventually developing into more mature AJs^41^. Our study revealed that apical localization of afadin, mediated by IDRs, triggers the formation of linear junctions on the apical side of epithelial cells. Interestingly, the PDZ domain of afadin was not essential for apical localization, suggesting that nectins do not determine the localization of afadin; rather, afadin define AJ localization of nectins. Moreover, EGFP- afadinΔC+FUSn, in which the IDR of afadin was replaced with an FUS-derived IDR, was also localized to the apical AJ, but its distribution within the AJC was different. These results indicate that the IDR is responsible for the distribution of afadin within the AJC.

How does afadin localize to the apical AJ. Noordstra et al. recently reported that antiparallel cortical flows at nascent AJs exert tension on the α-catenin actin-binding domain, causing it to increase adhesion growth^42^. Similarly, Schwayer et al. observed that non-functional ZO-1 clusters accumulate at junctions through retrograde actomyosin flow^17^. These cortical actin flow may also play a role in the apical accumulation of afadin.

However, EGFP-AF_C, a fragment of afadin’s IDR of afadin containing the F- actin-binding (FAB) domain, rarely localizes to apical junctions **(Fig. S4E)**. Additionally, afadinΔC, which retains the FAB domain but lacks the IDR, also show reduced accumulation at apical junctions **(Fig. 2A)**. These findings suggest that binding to the actin cytoskeleton alone is insufficient for the apical localization of afadin. Interestingly, the fact that afadin’s IDR localized to the apical junction even when replaced with an IDR from another molecule suggests that afadin’s IDR functions to facilitate the efficient clustering of molecules. Moreover, the addition of either the PDZ or RA domains to the IDR and FAB domains was also shown to localize these afadin fragments to apical junctions **(Fig. 6A)**. These results indicate that while the loss of a single domain in afadin does not significantly affect its localization to the apical AJ **(Fig. S1D)**, multivalent interactions via multiple domains, including IDRs, are essential for proper localization to AJ.

Through these multivalent interactions involving multiple domains of afadin with adhesion molecules, signaling molecules, and the actin cytoskeleton may create a local high-concentration environment. This, in turn, could trigger assembly via multiple weak hydrophobic interactions mediated by the IDR, leading to the formation of adhesive structures via LLPS. For instance, afadin’s interaction with nectin near the membrane, as well as with the actin cytoskeleton, could induce droplet-like structures through localized protein condensation. These condensates might concentrate other adhesion molecules such as cadherins and catenins, enabling their proper localization to apical junctions via interactions with the actin cytoskeleton.

This apical accumulation may involve retrograde flow, mediated by the actin cytoskeleton as mentioned earlier. During the initial phase of junction formation, afadin and adhesion molecules, which are distributed at relatively low concentrations in the lateral cell-cell adhesion, become linearly concentrated at the apical side as they move with retrograde flow. This mechanism may drive the formation of linear AJs at the apical side and promote cell height increase, thereby playing a role in establishing or maintaining cell polarity.

### IDR-dependent molecular segregation between afadin and ZO-1

In the present study, we showed that condensates of endogenous afadin and ZO-1 segregated in the juxta-membrane region in Fig.4G. Furthermore, in the experiment of co-expression of afadin and ZO-1, these molecules were not evenly distributed in the condensates **(Fig. 5F)**. These results suggest that molecular segregation between afadin and ZO-1 is possibly involved in the sorting of two distinct junctional domains during epithelial polarization. Previous studies have shown that in mature epithelial cells, afadin and ZO-1 localize to distinct adhesion structures, whereas in initial phase of epithelial polarization, they are found in the primordial AJs^30,35,43^. In the present study, we demonstrated that endogenous afadin and ZO-1 segregate into distinct condensates near the cell membrane **(Fig. 4G)**. Additionally, in co-expression experiments with afadin and ZO-1, these molecules did not distribute uniformly within the condensates **(Fig. 5F)**. These findings suggest that molecular segregation between afadin and ZO-1 may contribute to the sorting of distinct junctional domains during epithelial polarization. In afadin-KO cells, no abnormalities were observed in the localization of ZO-1 or Par-3, and epithelial barrier function was unaffected^14^. This indicates that AJ and TJ formation may be regulated independently, at least in cultured epithelial cells. This supports the idea that ZO-1 serves as the organizer of TJs, while afadin functions as the organizer of AJs. Recently, Mangeol et al. reported that the zonula adherens (ZA) is composed of two distinct adhesive belts: an apical nectin/afadin complex and a basal cadherin/catenin complex^44^. The structural segregation of TJs and AJs within the AJC is likely driven by the autonomous separation of afadin and ZO-1. Since afadin can form condensates through LLPS, while αE-catenin lacks this ability, the gradual separation of condensates containing both nectin/afadin and cadherin/catenin complexes may contribute to the layered structure of linear AJs.

## EXPERIMENTAL MODEL AND SUBJECT DETAILS

MDCK-II^45^ and HEK293^46^ cells were grown in a Dulbecco’s Modified Eagle Medium: Nutrient Mixture F-12 Ham (Gibco) supplemented with 10% fetal bovine serum (FBS) (MP Biomedicals) (DH10) at 37°C with 5% CO2, these cells were kindly gifted by Dr. Masatoshi Takeichi (RIKEN Center for Biosystems Dynamics Research, Kobe, Japan). Generation of afadin KO, ZO-1/2 dKO, and αE-catenin KO MDCK cells was previously described^14^.

## METHOD DETAILS

### Three-dimensional Cell culture

For three-dimensional (3D) culture, MDCK cells were embedded in bovine dermis atelocollagen gel (Koken). Briefly, atelocollagen gel and cell suspension prepared in DH10 and 20 mM HEPES (Gibco) were mixed at a 4:6 ratio on ice, and 200 μl with 4.5×10^4^ cells in the medium was plated on 13-mm-diameter cover glass (Matsunami Glass) in 24-well plate (Nunc) or 35-mm-diameter glass bottom dishes (Matsunami Glass). The gel was allowed to form at 37°C under 5% CO2 for 30 min, and 500 μl of warm DH10 was added into the plate or dish.

### Expression vectors

pCA-ZO-1-HA, pCAH-ZO-1-EGFP, pCANw-ZO-1-EGFP, pCAH-nectin-1, and pCAH-nectin-3 were previously described^14,47–49^. The following expression vectors were used, pCANw-sal-EGFP, pCAH, and pCAB^50^, and the following vectors were modified from the aforementioned vectors, pCANw-EGFP-C3, pCANw-mScarlet-C3, and pCANw- sal-mScarlet. mScarlet cDNA (without SalI, NotI, AhdI, and SacII restrict enzymic sites) was synthesized at Genewiz^51^. The following afadin mutants and chimeras were used by PCR amplification and Gibson assembly: The full-length cDNA for rat afadin (amino acids 1-1828) and cDNAs for afadinΔRA (amino acids 351-1828^52^), afadinΔPDZ (amino acids 1-1015 and 1122-1828^53^), AF_N-PDZ (amino acids 1-1113^30^, afadinΔC (amino acids 1-1113 and 1655-1828), afadinΔPR1-2 (amino acids 1-1218 and 1400-1828^30^), afadinΔCC (amino acids 1-1399 and 1461-1828^29^), s-afadin^54^, AF_1114-1654 (amino acids 1114-1654), afadinΔC+CC (amino acids 1-1113, 1400-1466, and 1655-1828), afadinΔC+FUSn (amino acids 1-1113, human FUS_1-214^39^, and 1655-1828), AF_PDZ (amino acids 990-1117), AF_1114-1654 (amino acids 1114-1654), AF_C (amino acids 1114-1828), AF_RA-1114-1654 (amino acids 1-350 and 1114-1654), AF_PDZ-1654 (amino acids 1016-1654), AF_DIL-PDZ-1654 (amino acids 647-1654), AF_RA-C (amino acids 1-350 and 1114-1828), AF_PDZ-C (amino acids 1016-1828), AF_DIL- PDZ-C (amino acids 647-1828), AF_RA-ZO-1_C (amino acids 1-350 and mouse ZO- 1_C_889-1755^16^), AF_PDZ-ZO-1_C (amino acids 1016-1122 and mouse ZO-1_C_889-1755), AF_DIL-PDZ-ZO-1_C (amino acids 647-1122 and mouse ZO-1_C_889-1755), AF_RA-ZO-1_PSG-U6-C (amino acids 1-350 and mouse ZO-1_PSG-U6-C_516- 1755^16^), AF_PDZ-ZO-1_PSG-U6-C (amino acids 1016-1122 and mouse ZO-1_PSG- U6-C_516-1755), AF_DIL-PDZ-ZO-1_PSG-U6-C (amino acids 647-1122 and mouse ZO-1_PSG-U6-C_516-1755), AF_RA-FUSn (amino acids 1-350 and human FUS_1- 214^39^), AF_PDZ-FUSn (amino acids 1016-1122 and human FUS_1-214), AF_DIL- PDZ-FUSn (amino acids 647-1122 and human FUS_1-214), afadinΔC+FUSn (amino acids 1-1113, human FUS_1-214, and 1655-1828) and afadinΔC+PSG (amino acids 1-1113, mouse ZO-1_PSG_516-810^40^, and 1655-1828). Afadin cDNAs were also subcloned into pCANw-EGFP-C3, pCANw-mScarlet-C3, pCAH-EGFP-C3, and pCAH-mScarlet-C3 using KOD One PCR Master Mix (#KMM-101X5, TOYOBO), Q5 High-Fidelity 2X Master Mix (New England Biolabs), and NEBuilder HiFi DNA Assembly Master Mix (New England Biolabs), according to the manufacturer’s instructions. Mouse αE-catenin cDNA^22^ was subcloned into pCANw-sal-EGFP and pCANw-sal-mScarlet by digestion and ligation. The following ZO-1 chimeras were used by PCR amplification and Gibson assembly: ZO-1ΔC-AF_C (amino acids 1-888^16^ and rat afadin_1114-1828) and ZO-1ΔC-FUSn (amino acids 1-888 and human FUS_1-214). ZO-1 cDNAs were also subcloned into pCANw-sal-EGFP. See also key resource tables and Figs. S1A&S4A&S5.

### DNA transfection

HEK293 cells were transfected with various combinations of expression vectors in 10- cm dishes. Transfection were performed with PEI MAX (Polysciences), Lipofectamine 3000 (Invitrogen), HilyMax (Dojindo), or Effectene (QIAGEN) according to the manufacturer’s instructions. Afadin KO, αE-catenin KO, and ZO-1/2 dKO MDCK cells were transfected with expression vectors, respectively. Transfection of expression vectors was performed with PEI MAX, Lipofectamine 3000, HilyMax, or Effectene according to the manufacturer’s instruction.

### Generation of stable transfectants

The transfected MDCK cells were selected using 400 μg/ml G418 disulfate (NacalaiTesque) or 200 μg/ml hygromycin B Gold (Invivogen), the mixed clones with the lowest EGFP expression were collected by a FACS (SH800Z, SONY). These multiple clones were maintained in the presence of G418 disulfate or hygromycin Gold B.

### Antibodies and Reagents

The following primary antibodies (Ab) were used: anti-l-afadin mouse monoclonal Ab (mAb)(clone 3, OriGene; 1:50 dilution for immunofluorescence, 1:3000 dilution for western blotting), anti-α-catenin rabbit polyclonal Ab (pAb) (Sigma-Aldrich; 1:400 dilution for immunofluorescence, 1:1000 dilution for western blotting), anti-Par-3 rabbit pAb (Sigma-Aldrich; 1:100 dilution for immunofluorescence), anti-β-catenin mouse mAb (clone 5H10^55^, Sigma-Aldrich; 1:1000 dilution for immunofluorescence), anti-β- catenin rabbit pAb (Sigma-Aldrich; 1:800 for immunofluorescence), anti-ZO-1 rat mAb (clone R40.76^56^, Santa Cruz Biotechnology; 1:50 dilution for immunofluorescence, 1:1000 dilution for western blotting), anti-HA rat mAb (clone 3F10, Roche; 1:4000 dilution for western blotting), anti-α-tubulin mouse mAb (clone DM1A^57^, Sigma- Aldrich, 1:5000 dilution for western blotting), anti-GFP rat mAb (clone GF090R, NacalaiTesque; 2.5 μg/reaction for co-immunoprecipitation).

For immunofluorescence, primary Abs were visualized with secondary goat antibodies conjugated with Alexa Fluor plus 405, Alexa Fluor 488, Alexa Fluor 647 (Invitrogen; Jackson Immunoresearch Laboratories; 1:400 dilution), and Cy3 (Sigma-Aldrich; 1:400 dilution).

For nuclei and F-actin staining were visualized with 4’,6-diamidino-2- phenylindole (DAPI) (Dojindo; 1:1000 dilution) and phalloidin Alexa Fluor plus 405- conjugated (Invitrogen; 1:400 dilution), phalloidin ATTO 565-conjugated (Sigma-Aldrich; 1:200 dilution), and phalloidin Alexa Fluor 647-conjugated (Invitrogen, 1:100 dilution). For western blotting, primary Abs were detected with secondary rabbit, mouse, and rat pAbs horseradish peroxidase-conjugated (HRP-Ab) (Jackson Immunoresearch Laboratories; Invitrogen; 1:5000 dilution).

The following inhibitors with dimethyl sulfoxide (DMSO) (FUJIFILM Wako Pure Chemical Corporation) were used: 1,6-hexanediol (1,6-HD)^31,32^ (NacalaiTesque), Y-27632^28^ (FUJIFILM Wako Pure Chemical Corporation), and latrunculin B (LatB) (Tocris Bioscience).

### Immunofluorescence

Cultured cells for 24 hours of incubation were fixed with 4% paraformaldehyde (PFA) (Merck) in Hank’s balanced salt solution containing 1 mM Ca^2+^ and Mg^2+^ (Gibco), containing 10 mM HEPES (Gibco) (HBSS CM+) at 37°C for 15 min. After fixation, the samples were washed three times with Tris-buffered saline, pH 7.6 containing 1 mM Ca^2+^ (NacalaiTesque) and 0.005% Tween20 (NacalaiTesque) (TBS-C) and permeabilized with a 0.25% Triton X-100 (NacalaiTesque) in TBS-C at room temperature (RT) for 5 min. The samples were then blocked with 5% normal goat serum (Sigma-Aldrich) or 5% skim milk (Morinaga milk industry) in TBS-C at RT for 30 min, followed by incubation with primary antibodies in 2.5% normal goat serum or 2.5% skim milk in TBS-C or in “Can Get Signal” immunoreaction enhancer solution B (TOYOBO) at RT for 2 h or 4°C overnight. The samples were washed three times with TBS-C, then incubated with secondary Abs and counterstain dyes in 2.5% normal goat serum or 2.5% skim milk in TBS-C at RT for an hour. These samples were washed three times with TBS-C and mounted on a slide glass with CC/Mount (Diagnostic Biosystems) or Fluorosave reagent (Millipore). For super-resolution imaging, samples were mounted with RapiClear 1.52 reagent (SunJin Lab) and iSpacer (SunJin Lab).

3D culture cells for 8 days were incubated with collagenase type VII (Sigma- Aldrich) at RT for 15 min. The cells were fixed with 2% PFA in HBSS CM+ at 37°C for 30 min. After another three rinses with TBS-C, the cells were blocked with 0.5% Triton X-100 and 5% normal goat serum in TBS-C at RT for 30 min. The primary Abs were diluted in “Can Get Signal” immunoreaction enhancer solution B and incubated with the gels at 4°C overnight with gentle rocking. The gels were washed with TBS-C five times and incubated with the secondary Abs at 4°C overnight with gentle rocking. After five washes with TBS-C, the samples were mounted in CC/Mount. A tall sample was mounted into the bank of a microscope slide by hollowing out the inside of vinyl tape.

Images were obtained with confocal laser scanning microscopes (AX-R, Nikon Corporation; LSM700, LSM900 with Airyscan2, and LSM980 with Airyscan2, Carl Zeiss Microscopy; FV1000, OLYMPUS Life Science) equipped with a CFI Plan Apochromat Lambda D 60x/1.42 (Nikon Corporation), a LD LCI Plan-Apochromat 40x/1.2 (Carl Zeiss Microscopy), C-Apochromat 40x/1.2 W Korr (Carl Zeiss Microscopy), Plan-Apochromat 63x/1.4 (Carl Zeiss Microscopy), a UPLSAPO 20x/0.75 (OLYMPUS Life Science), and a PLAPON60XOSC2 60x/1.4 (OLYMPUS Life Science) lens. These images were processed and analyzed with ZEN 3.9 software into “bioformat import” function (Carl Zeiss Microscopy). Line fluorescence intensities were scanned along arrows and normalized per 6 pixels using the “Profile View” function. Cell height was measured using the “Ortho View” function of nuclei and β-catenin signals. Cell height of Cyst was measured using the β-catenin Z-projection signal. The tilting extent was measured as referenced in the previous report^5^. The degree of tilting of LCs was quantified as follows: LC area was defined as EGFP-afadins or β-catenin signals. Then, the width of the border between the LC and AJC areas was measured. Subsequently, the LC area was divided by the width of the border to obtain a mean LC area per AJC unit (1 μm). This value was defined as the “tilting extent.” Condensate number of the cell was measured by orthogonal projection images. To compare the colocalization of EGFP-afadin and β-catenin in the cyst, we used the “colocalization colormap” plugin of Fiji^58,59^. We cropped three images of 50×50 pixels of apical areas, and quantified afadin and β-catenin signals, according to the developer’s instruction. Statistical tests and graphs were performed using Kyplot 6.0.2 software^60^.

### Western blotting

The samples separated by SDS-PAGE were transferred to polyvinylidene difluoride membranes (Millipore). After the membranes were blocked with Block ACE (KAC Co., Ltd.) in Tris-buffered saline, pH 7.6 containing 0.05% Tween 20 (TBS-T), they were incubated with the indicated Abs at 4°C overnight. After the membranes were washed with TBS-T three times, they were incubated with HRP-Abs. The membranes were then washed with TBS-T three times, and signals for the proteins were detected using Chemi-Lumi One L (NacalaiTesque) and a chemiluminescent image analyzer (LAS-4000, FUJIFILM Wako Pure Chemical Corporation).

### Co-immunoprecipitation

The transfected HEK293 cells were washed with phosphate-buffered saline and then suspended in 1 ml of Buffer A (20 mM Tris-HCl, pH 7.4, 150 mM NaCl, 1% Triton X- 100, 1 mM EDTA, 1 mM Na3VO4, and protease inhibitor cocktail for use with mammalian cell and tissue extracts (NacalaiTesque) on ice. The cell extract was obtained by centrifugation at 15,000 rpm at 4°C for 15 min. Protein loadings were normalized by protein assay BCA kit (NacalaiTesque). 500 μg of cell extracts was then incubated with 2.5 μg of anti-GFP mAb at 4°C overnight. The reactants were incubated with dynabeads protein G slurry (Invitrogen) at RT for an hour. After supernatant discard, the beads were extensively washed with Buffer A, and the bound proteins were eluted by boiling the beads in 55 μl of 6× SDS sample buffer (62.5 mM Tris-HCl, pH 6.8, 2% SDS, 0.1 M dithiothreitol, 10% glycerol, and 0.01% bromophenol blue) for 5 min. Each 10 μl of the samples was then subjected to SDS-PAGE, followed by western blotting with the anti- afadin, -HA, and -α-catenin.

### Live cell imaging

Live imaging of MDCK cells was performed using a confocal laser scanning microscope (FV1000, OLYMPUS Life Science) equipped with a standard heating stage top incubator (INUB-ZILCS, TOKAI HIT) at 37°C under 5% CO2. The cells were seeded onto a 35-mm-diameter glass bottom dish (Matsunami Glass) and observed with a UPLSAPO30XS, or a PLAPON60XOSC2 lens. Before experiments, the cell media were replaced with 3 ml of FluoroBrite DMEM (Gibco) supplemented with 10% FBS and 10 mM HEPES. These images were processed and analyzed with ZEN software. 3D video rendered by using the “3D View” function. Line fluorescence intensities were scanned along arrows and normalized per 3 pixels using the “Profile View” function. Line graphs were made by Kyplot software.

### Fluorescence recovery after photobleaching (FRAP)

FRAP experiments in cells were conducted using an LSM980 with Airyscan2, equipped with a C-Apochromat 40x/1.2 W Korr objective (Carl Zeiss Microscopy). A region of interest (ROI) was bleached using a 488 nm or 561 nm laser at 10 mW power with a pixel dwell time of 100 µs. Pre-bleach and post-bleach images were acquired using the 488 nm or 561 nm laser. Fluorescence recovery of EGFP or mScarlet was monitored for 1-2 min with a time resolution of 30 or 180 ms. Recovery data was background corrected and normalized to the ROI intensity prior to bleaching. A reference ROI outside the bleached area was processed in the same way.

### Fluorescence correlation spectroscopy (FCS)

FCS experiments in cells were performed using an LSM980 with an Airyscan2 and a C-Apochromat 40x/1.2 W Korr objective lens and analyzed with ZEN Dynamics Profiler for FCS software (Carl Zeiss Microscopy). mScarlet was excited with a 561 nm laser and the fluorescence fluctuations were recorded with a time resolution of 1.2 μs for 20 s. Autocorrelation of the photon traces was performed in ZEN Dynamics Profiler for FCS software using detrending filter (Carl Zeiss Microscopy), and the resulting correlation curves were fitted according to the 2-component 3D diffusion model.

### Aliphatic alcohols treatment on cells

HEK293 cells, which express EGFP, EGFP-afadin, or ZO-1-EGFP, were incubated in DH10 supplemented with 5-10% 1,6-HD (weight/volume, Sigma-Aldrich) for 5 min and fixed with 4% PFA at 37°C for 30 min. Alternatively, MDCK cells were incubated in DH10 supplemented with 0.5% 1,6-HD for 60 min and then fixed with 4% PFA in HBSS CM+ at 37°C for 30 min.

### Disassembly of the F-actin polymerization on MDCK cells

Confluent monolayers of afadin KO MDCK cells expressing EGFP-afadin or EGFP- afadinΔC, were incubated in FluoroBrite DMEM supplemented with 1 μM LatB. Live cell imaging was acquired every 3 min for a duration of over 180 min after LatB addition.

### Disassembly of the F-actin polymerization on HEK293 cells

HEK293 cells, which transiently expressing EGFP-afadin, were incubated in DH10 supplemented with DMSO or 1 μM LatB for 90 min. Cells were fixed with 4% PFA in HBSS CM+ at 37°C for 30 min. The samples were washed three times with TBS-C and then incubated with DAPI in TBS-C at RT for one hour. EGFP signals were measured for number, area, long axis, and circularity. Circularity was calculated as: *Circularity =*

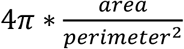

A circularity value of 1 reflects a perfect circle, while values close to 0 indicate more elongated shapes.

### Client partitioning assays in HEK293 cells

HEK293 cells were transfected with EGFP-afadin or ZO-1-EGFP and clients were tagged with mScarlet. The clients were mScarlet, mScarlet-afadin, mScarlet-afadinΔC, and αE-catenin-mScarlet. Cells were fixed with 4% PFA in HBSS CM+ at 37°C for 30 minutes. The samples were washed three times with TBS-C and then incubated with DAPI in TBS-C at RT for one hour. To quantify the partition co-efficient of the client, we determined the region of condensate using EGFP signals. We then measured the mean intensity of the client condensate/cytoplasm in the condensate region and subtracted the extracellular background from these values. The partition co-efficient was calculated as:

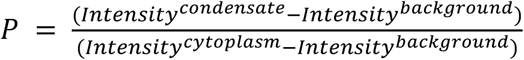

Intensity^cytoplasm^ and intensity^background^ averaged over 10 μm^2^. Graphs were visualized as log10P. A log10P value of 0 reflects equal partitioning of condensate and cytoplasm, while values higher than 0 indicate more clients in the condensate.

### Mosaic forming assay

2-well culture inserts (ibidi) were attached to glass-bottomed dishes. WT or afadin KO MDCK cells expressing nectin-1 and EGFP or nectin-3 and mScarlet, were seeded into each well of the culture insert. When the cells formed confluent monolayers after 2-3 days, the culture inserts were detached with forceps. The cells were then cultured for another 2-3 days. DMEM without Ca^2+^ (Gibco) was incubated for 2 h. To induce cell migration, a vertical scratch as made through each cell colony using a pipette tip. Cells were cultured in DH10 for 2 days. The cells were fixed with 4% PFA in HBSS CM+ at 37°C for 30 min. The samples were washed three times with TBS-C, then incubated with DAPI in TBS-C at RT for one hour. The number of isolated single cells appearing in the different colored colonies of the other cells was counted for each combination.

## Quantification and statistical analysis

Quantification was represented by the mean ± standard deviation. We performed the Mann-Whitney *U*-test, Steel’s multiple comparison test, and Steel and Dwass’s multiple comparison test for the non-parametric statistical analyses, as indicated in the figure legends, using Kyplot software. P values of less than 0.05 were regarded as statistically significant.

## Data and code availability

All data supporting the current study are available from the corresponding author upon request.

## Acknowledgments

The authors are sincerely grateful to Drs. Suzuki, A., Nitta, R., Otani, J., Katsunuma, S., and Ueyama, T (Kobe University). for discussions; Nikon imaging center of Osaka University, Mimori-Kiyosue, Y (RIKEN BDR), Lin, CY (SunJin Lab Co.), Kamei, Y (National Institute for Basic Biology), Ishidate, F (Kyoto University), Sato, Y (Carl Zeiss, Japan), for technical assistance. This work was supported by JST PRESTO Grant Number JPMJPR1946 (H.T.); by JST SPRING Grant Number JPMJSP2148 (S.K.); by KAKENHI Grants Numbers JP19K03634(H.T.), JP20H01823(H.T.), JP22K19331(H.T.), JP22H04926(ABiS), and JP24H00188(H.T.); by Grant-in-Aid for Transformative Research Areas - Platforms for Advanced Technologies and Research Resources “ABiS (Advanced Bioimaging Support)”; by a grant from the Takeda Science Foundation (H.T.); and by Iue Memorial Foundation (S.K.).

The authors declare no conflict of interest.

## Abbreviations List

Abbreviations used in this paper: AJ, adherens junction; TJ, tight junction; AJC, apical junctional complex; IDR, intrinsically disordered region; EGFP, enhanced green fluorescent protein; FACS, fluorescence-activated cell sorting; LC, lateral contact; LLPS, liquid-liquid phase separation; ROI, region of interest; ZA, zonula adherens; h, hours; min, minutes; s, seconds.

## Supplemental information

Figure S1. Characterization of afadin KO, ZO-1/2 dKO, and αE-catenin KO MDCK cells, related to figure 1.

Figure S2. Localizations of afadin mutants upon expression in afadin KO, ZO-1/2 KO, or αE-catenin KO cells, related to figure 2.

Figure S3. Junctional dynamics of afadin mutants upon expression in afadin KO, ZO-1/2 KO, or αE-catenin KO cells, related to figure 3.

Figure S4. Schematic diagram of afadin IDR mutants and the localization of afadin IDR fragments, related to figure 4.

Figure S5. Schematic diagram of systematic afadin extended fragments and ZO-1- Afadin chimeras, related to figure 6&7.

Video S1 Junction formation of nectin-1 and AF_PDZ-EGFP expressing WT MDCK cells, related to figure 2.

Video S2 Junction formation of nectin-1 and AF_PDZ-EGFP expressing afadin KO cells, related to figure 2.

Video S3 Junction formation of EGFP-afadin expressing afadin KO cells, related to figure 3.

Video S4 Junction formation of EGFP-afadinΔC expressing afadin KO cells, related to figure 3.

Video S5 Junction formation of nectin-1 and EGFP-afadin expressing afadin KO cells, related to figure 3.

Video S6 Junction formation of nectin-1 and EGFP-afadinΔC expressing afadin KO cells, related to figure 3.

Video S7 Junction formation of EGFP-afadinΔPR1-2 expressing afadin KO cells, related to figure S3.

Video S8 Junction formation of EGFP-afadinΔCC expressing afadin KO cells, related to figure S3.

Video S9 Junction formation of EGFP-afadinΔC+CC expressing afadin KO cells, related to figure S3.

Video S10 Junction formation of EGFP-afadin expressing ZO-1/2 dKO cells, related to figure S3.

Video S11 Junction formation of EGFP-afadinΔC expressing ZO-1/2 dKO cells, related to figure S3.

Video S12 Junction formation of EGFP-afadin expressing αE-catenin KO cells, related to figure S3.

Video S13 Junction formation of EGFP-afadinΔC expressing αE-catenin KO cells, related to figure S3.

Video S14 Condensates formation of EGFP-afadin, related to figure 4.

Video S15 Molecular dynamics of EGFP-afadin by the treatment of LatB, related to figure 4.

Video S16 Molecular dynamics of EGFP-afadinΔC by the treatment of LatB, related to figure 4.

Video S17 Recovery dynamics of an EGFP-afadin expressing HEK293 cells by FRAP, related to figure 5.

Video S18 Recovery dynamics of junctional mScarlet-afadin expressing ZO-1/2 dKO cells by the FRAP, related to figure 5.

Video S19 Recovery dynamics of junctional mScarlet-afadinΔC expressing ZO-1/2 dKO cells by the FRAP, related to figure 5.

## Supplemental information

## Video legends

**Video S1 Junction formation of nectin-1 and AF_PDZ-EGFP expressing WT MDCK cells, related to** figure 2. Apical views of the transfectants. Cells were cultured for 24 h. Time-lapse images of the junction formation. Cells were observed for 850 min at 10 min intervals. Scale bar, 10 μm.

Video S2 Junction formation of nectin-1 and AF_PDZ-EGFP expressing afadin KO cells, related to figure 2. Apical views of the transfectants. Cells were cultured for 24 h. Time-lapse images of the junction formation. Cells were observed for 850 min at 10 min intervals. Scale bar, 10 μm.

Video S3 Junction formation of EGFP-afadin expressing afadin KO cells, related to figure 3. Apical views of the transfectants. Cells were cultured for 24 h. Time-lapse images of the junction formation. Cells were observed for 250 min at 10 min intervals. Scale bar, 10 μm.

Video S4 Junction formation of EGFP-afadinΔC expressing afadin KO cells, related to figure 3. Apical views of the transfectants. Cells were cultured for 24 h. Time-lapse images of the junction formation. Cells were observed for 250 min at 10 min intervals. Scale bar, 10

Video S5 Junction formation of nectin-1 and EGFP-afadin expressing afadin KO cells, related to figure 3. Apical views of the transfectants. Cells were cultured for 24 h. Time-lapse images of the junction formation. Cells were observed for 250 min at 10 min intervals. Scale bar, 10 μm.

Video S6 Junction formation of nectin-1 and EGFP-afadinΔC expressing afadin KO cells, related to figure 3. Apical views of the transfectants. Cells were cultured for 24 h. Time-lapse images of the junction formation. Cells were observed for 250 min at 10 min intervals. Scale bar, 10μm.

Video S7 Junction formation of EGFP-afadinΔPR1-2 expressing afadin KO cells, related to figure S3. Apical views of the transfectants. Cells were cultured for 24 h. Time-lapse images of the junction formation. Cells were observed for 250 min at 10 min intervals. Scale bar, 10 μm.

Video S8 Junction formation of EGFP-afadinΔCC expressing afadin KO cells, related to figure S3. Apical views of the transfectants. Cells were cultured for 24 h. Time-lapse images of the junction formation. Cells were observed for 250 min at 10 min intervals. Scale bar, 10

Video S9 Junction formation of EGFP-afadinΔC+CC expressing afadin KO cells, related to figure S3. Apical views of the transfectants. Cells were cultured for 24 h. Time-lapse images of the junction formation. Cells were observed for 250 min at 10 min intervals. Scale bar, 10 μm.

Video S10 Junction formation of EGFP-afadin expressing ZO-1/2 dKO cells, related to figure S3. Apical views of the transfectants. Cells were cultured for 24 h. Time-lapse images of the junction formation. Cells were observed for 250 min at 10 min intervals. Scale bar, 10 μm.

Video S11 Junction formation of EGFP-afadinΔC expressing ZO-1/2 dKO cells, related to figure S3. Apical views of the transfectants. Cells were cultured for 24 h. Time-lapse images of the junction formation. Cells were observed for 250 min at 10 min intervals. Scale bar, 10 μm.

Video S12 Junction formation of EGFP-afadin expressing αE-catenin KO cells, related to figure S3. Apical views of the transfectants. Cells were cultured for 24 h. Time-lapse images of the junction formation. Cells were observed for 250 min at 10 min intervals. Scale bar, 10

Video S13 Junction formation of EGFP-afadinΔC expressing αE-catenin KO cells, related to figure S3. Apical views of the transfectants. Cells were cultured for 24 h. Time-lapse images of the junction formation. Cells were observed for 250 min at 10 min intervals. Scale bar, 10 μm.

Video S14 Condensates formation of EGFP-afadin, related to figure 4. 3D views of the transfectants. The cell was transfected transiently after 3 days. This cell was observed for 200 min at 1 min intervals. The upper side of the video shows the apical side of the cell, and the lower side shows the basal side. Grid scales indicate per 2 μm.

Video S15 Molecular dynamics of EGFP-afadin by the treatment of LatB, related to figure 4. Apical views of the transfectants. These confluent cells were observed for 180 min at 3 min intervals in the presence of 1 μM LatB. The look-up table represents the strength of the EGFP signal, with magenta indicating the maximum threshold and black indicating the minimum threshold. Scale bar, 2 μm.

Video S16 Molecular dynamics of EGFP-afadinΔC by the treatment of LatB, related to figure 4. Apical views of the transfectants. These confluent cells were observed for 180 min at 3 min intervals in the presence of 1 μM LatB. The look-up table represents the strength of the EGFP signal, with magenta indicating the maximum threshold and black indicating the minimum threshold. Scale bar, 2 μm.

Video S17 Recovery dynamics of an EGFP-afadin expressing HEK293 cells by FRAP, related to figure 5. The same Z slice views of the EGFP-afadin condensate. The cell was transfected transiently after 3 days. After the observation from 1.5 s, a region of interest (ROI) was bleached using a 488 nm laser at 10 mW power with a pixel dwell time of 100 μs.

Video S18 Recovery dynamics of junctional mScarlet-afadin expressing ZO-1/2 dKO cells by the FRAP, related to figure 5. The same Z slice views of junctional mScarlet-afadin. Cells were cultured for 3 days. After the observation from 9 s, a ROI was bleached using a 561 nm laser at 10 mW power with a pixel dwell time of 100 μs. This cell junction was observed for 45 s at 180 ms intervals. Scale bar, 2 μm.

**Video S19 Recovery dynamics of junctional mScarlet-afadinΔC expressing ZO-1/2 dKO cells by the FRAP, related to** figure 5. The same Z slice views of junctional mScarlet-afadinΔC. Cells were cultured for 3 days. After the observation from 9 s, a ROI was bleached using a 561 nm laser at 10 mW power with a pixel dwell time of 100 μs. This cell junction was observed for 45 s at 180 ms intervals. Scale bar, 2 μm.

---

**Figure S1.**
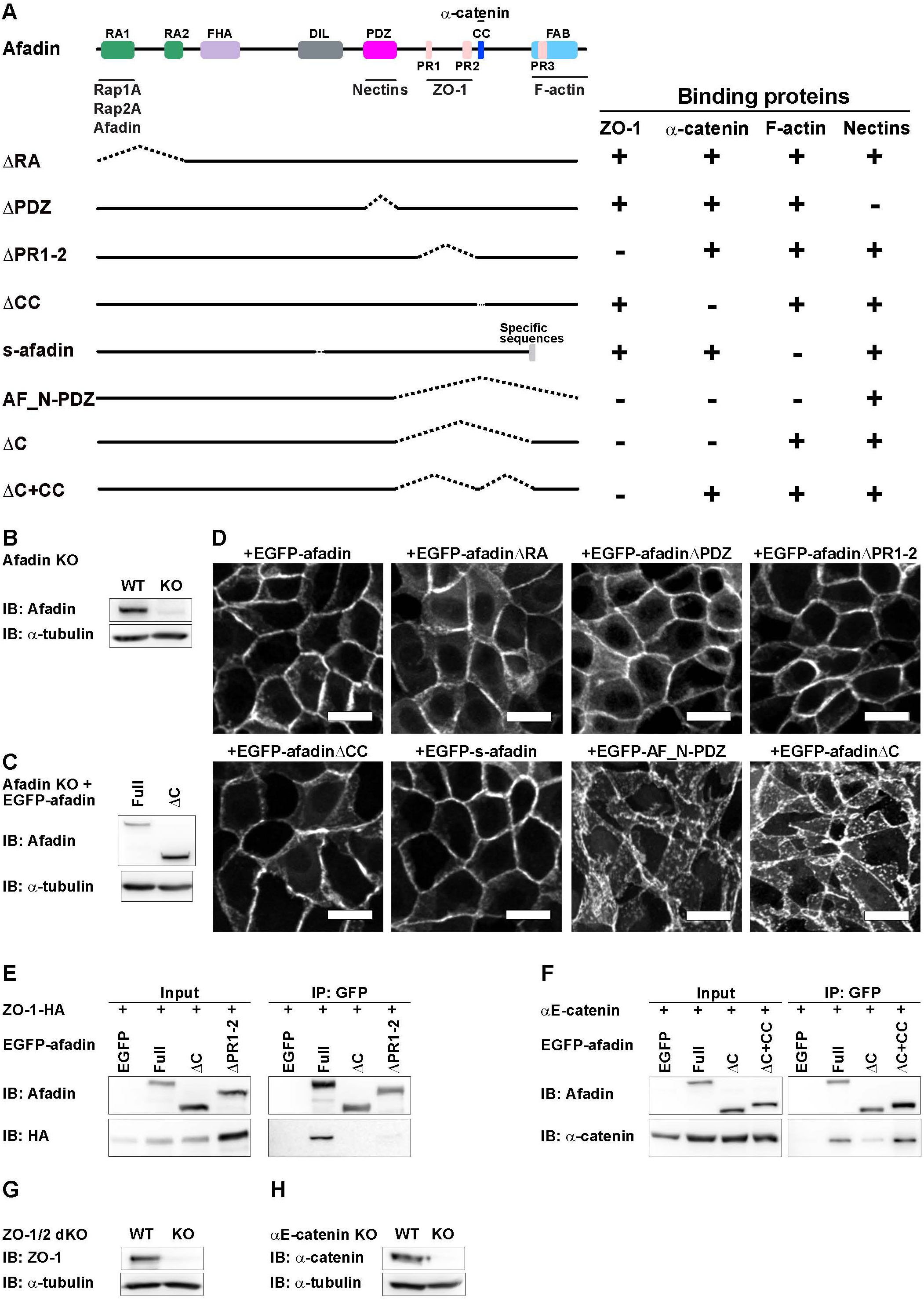
Characterization of afadin KO, ZO-1/2 dKO, and αE-catenin KO MDCK cells, related to. figure 1**. (A**) Schematic diagram of different afadin fragments. Afadin fragments were tagged with EGFP at the N-terminus. RA, Ras-associated domain; FHA, forkhead-associated domain; DIL, dilute domain; PDZ, PSD95-DLG1-ZO-1 domain; PR, proline-rich region; CC, coiled-coil domain; FAB, F-actin binding domain. +, positive for the interaction with ZO- 1, α-catenin, F-actin, or nectins; -, negative for the interaction with ZO-1, α-catenin, F-actin, or nectins. **(B)** Expression levels of afadin and α-tubulin in WT and afadin-KO cells. Cell extracts of WT and KO cells were separated by SDS-PAGE and subjected to western blotting using the indicated Abs. α-tubulin was immunoblotted as a loading control. IB, immunoblot. **(C)** Expression levels of EGFP-afadin and EGFP-afadinΔC in afadin-KO cells. The lowest expressing clones were collected by FACS. **(D)** Distribution of EGFP-afadin, EGFP-afadinΔRA, EGFP-afadinΔPDZ, EGFP-afadinΔPR1-2, EGFP- afadinΔCC, EGFP-s-afadin, EGFP-AF_N-PDZ, and EGFP-afadinΔC in afadin KO cells. The lowest expressing cells were observed by confocal microscopy. A representative image is shown of six independent experiments. Scale bars, 20 μm. **(E)** Co- immunoprecipitation of ZO-1-HA with the EGFP-afadins. Cell extracts of HEK293 cells transfected with various combinations of the indicated vectors were immunoprecipitated with the anti-GFP Ab. Immunoprecipitates and cell extracts were subjected to western blotting using the anti-afadin and anti-HA Abs. IP, immunoprecipitation. **(F)** Co- immunoprecipitation of α-catenin with the EGFP-afadins. Cell extracts of HEK293 cells transfected with various combinations of the indicated vectors were immunoprecipitated with the anti-GFP. Immunoprecipitates and cell extracts were subjected to western blotting using the anti-afadin and anti-α-catenin Abs. **(G)** Expression levels of ZO-1 and α-tubulin in WT and ZO-1/2 dKO cells. **(H)** Expression levels of α-catenin and α-tubulin in WT and αE-catenin KO cells.

**Figure S2.**
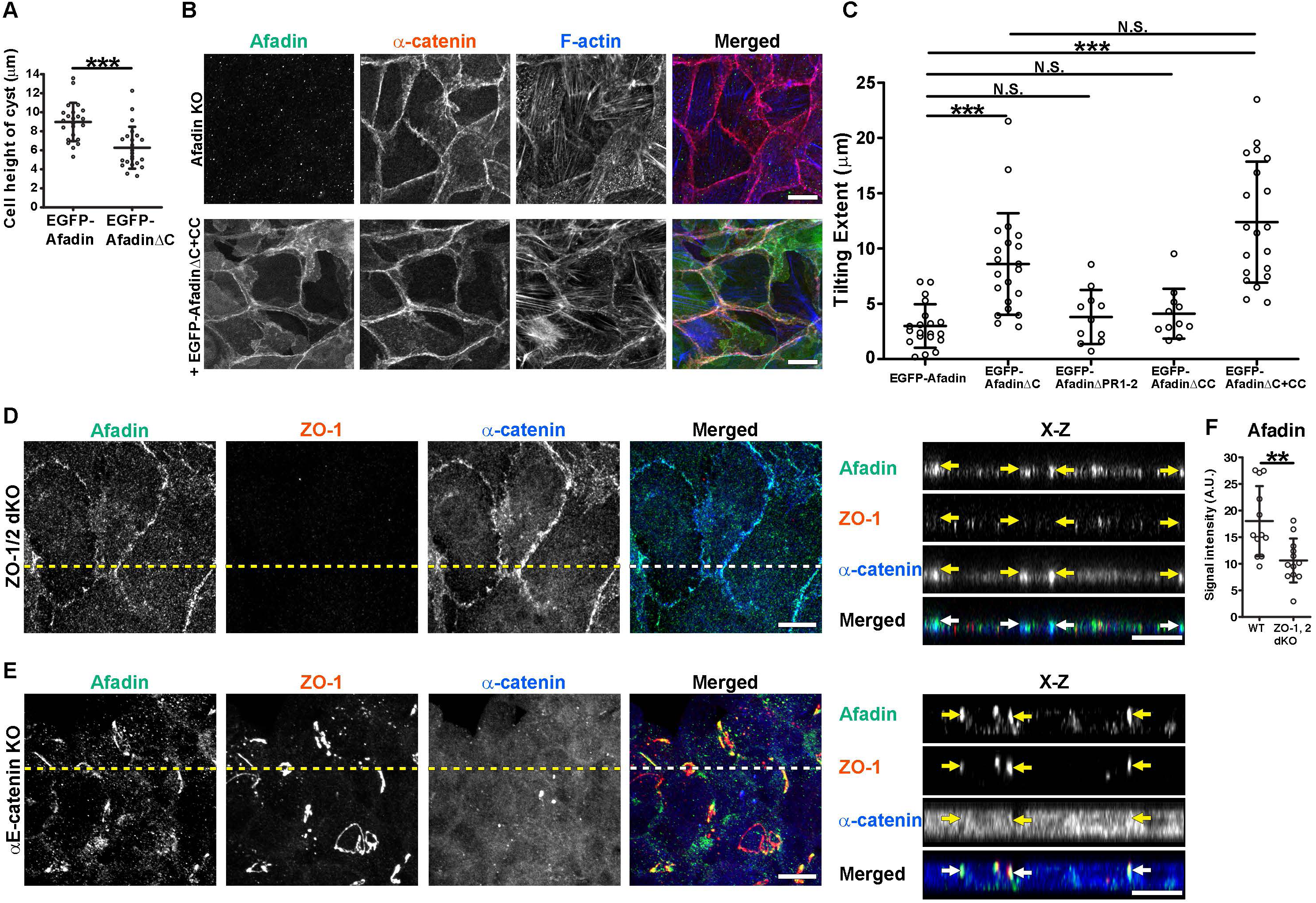
Localizations of afadin mutants upon expression in afadin KO, ZO-1/2 dKO, or αE-catenin KO cells, related to figure 2. **(A)** Quantifications of cell height of cyst in afadin-KO cells. Whiskers represent mean ± S.D. n = 23 for EGFP-afadin; n = 23 for EGFP-afadinΔC. Mann-Whitney *U*-test; ***, P ≤ 0.001. See also Figure 2B. **(B)** Distribution of endogenous afadin or EGFP- afadinΔC+CC (green), α-catenin (red), and F-actin (blue) in afadin-KO cells or EGFP- afadinΔC+CC-expressing afadin-KO cells; apical views of the transfectants cultured for 24 h and stained for the indicated molecules. Scale bars, 10 μm. Representative images are shown of six independent experiments. **(C)** Quantifications of tilting extent of LCs in afadin-KO cells. Whiskers represent mean ± S.D. Datasets for EGFP-afadin and EGFP- afadinΔC were equal to those in Figure 1C. Tilting extent; n = 20 for EGFP-afadin; n = 21 for EGFP-afadinΔC; n = 11 for EGFP-afadinΔPR1-2; n = 11 for EGFP-afadinΔCC; n = 20 for EGFP-afadinΔC+CC. Steel and Dwass’s multiple comparison test; ***, P ≤ 0.001; N.S., not significant. **(D)** Distribution of afadin (green), ZO-1 (red), and α-catenin (blue) in ZO-1/2 dKO cells; apical views of ZO-1/2 dKO cells cultured for 24 h and stained for the indicated molecules. Dashed lines indicate the position of vertical section. A representative image is shown of six independent experiments. Scale bar, 10 μm. (Right) Vertical sections of the left images. Arrows indicate the AJCs. Scale bar, 10 μm. **(E)** Distribution of afadin (green), ZO-1 (red), and α-catenin (blue) in αE-catenin KO cells; apical views of αE-catenin KO cells cultured for 24 h and stained for the indicated molecules. Dashed lines indicate the position of vertical sections. Representative images are shown of six independent experiments. Scale bar, 10 μm. (Right) Vertical sections of the left images. Arrows indicate the AJCs. Scale bar, 10 μm. **(F)** Quantifications of junctional anti-afadin fluorescence signals in WT and ZO-1/2 dKO cells. A.U., arbitrary unit. Whiskers represent mean ± S.D. n = 12 for WT; n = 12 for ZO-1/2 dKO. Mann- Whitney *U*-test; **, P ≤ 0.01.

**Figure S3.**
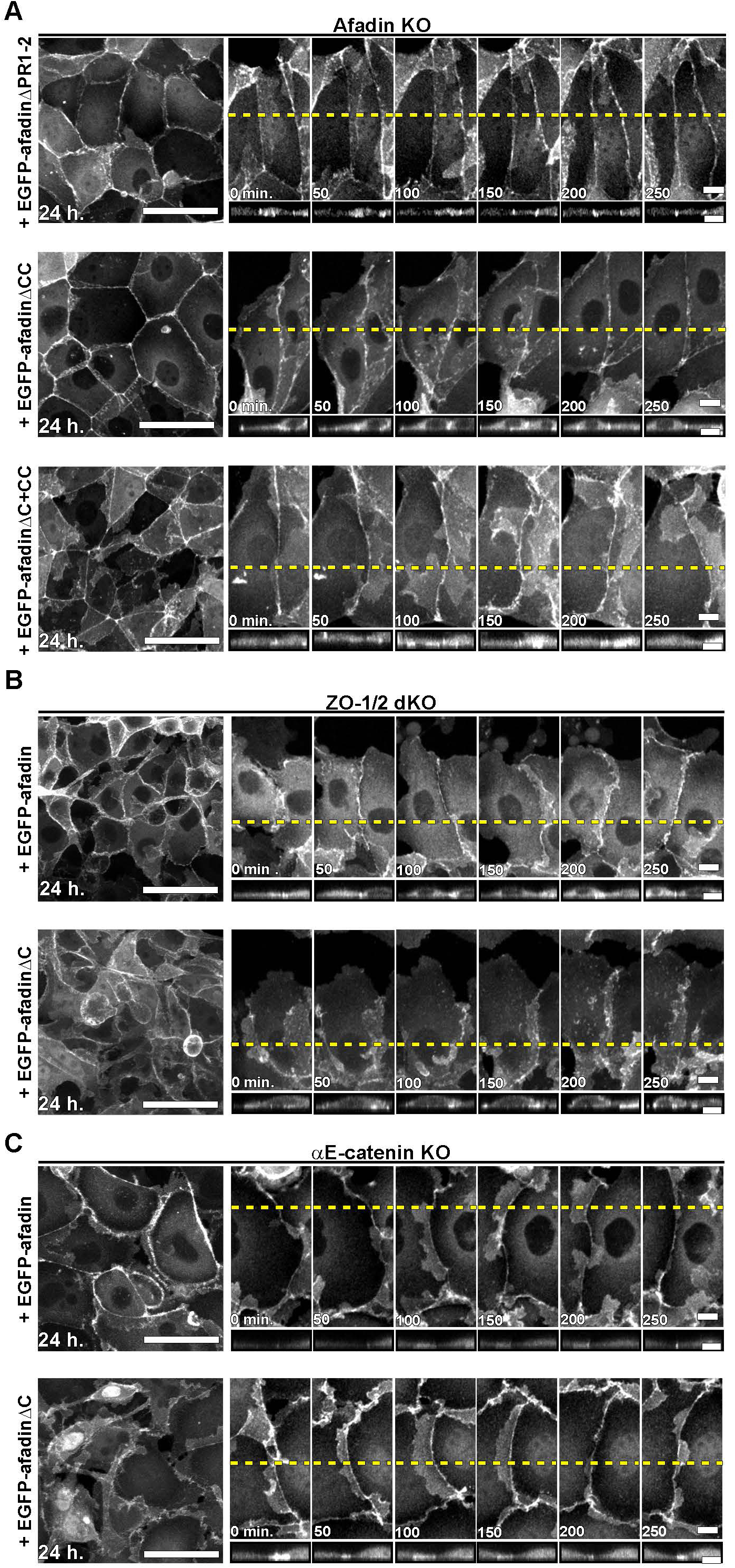
Junctional dynamics of afadin mutants upon expression in afadin KO, ZO-1/2 KO, or αE-catenin KO cells, related to figure 3. **(A)** Junction formation of EGFP-afadinΔPR1-2-, EGFP-afadinΔCC-, and EGFP- afadinΔC+CC-expressing afadin-KO cells. (Left) Apical views of the transfectant of the cells cultured for 24 h. Scale bars, 50 μm. (Right) Time-lapse images of the junction formation in cells observed for 250 min at 50 min intervals. See also Videos S6-S8. Scale bars, 10 μm. (Bottom) Vertical sections of the upper images. Scale bars, 10 μm. **(B)** Junction formation of EGFP-afadin- and EGFP-afadinΔC-expressing ZO-1/2-dKO cells. (Left) Apical views of the transfectant. These cells were cultured for 24 h. Scale bars, 50 μm. (Right) Time-lapse images of the junction formation. These cells were observed for 250 min at 50 min intervals. See also Videos S9&S10. Scale bars, 10 μm. (Bottom) Vertical sections of the upper images. Scale bars, 10 μm. **(C)** Junction formation of EGFP- afadin- and EGFP-afadinΔC-expressing αE-catenin-KO cells. (Left) Apical views of the transfectant. These cells were cultured for 24 h. Scale bars, 50 μm. (Right) Time-lapse images of the junction formation. These cells were observed for 250 min at 50 min intervals. See also Videos S11&S12. Scale bars, 10 μm. (Bottom) Vertical sections of the upper images. Scale bars, 10 μm.

**Figure S4.**
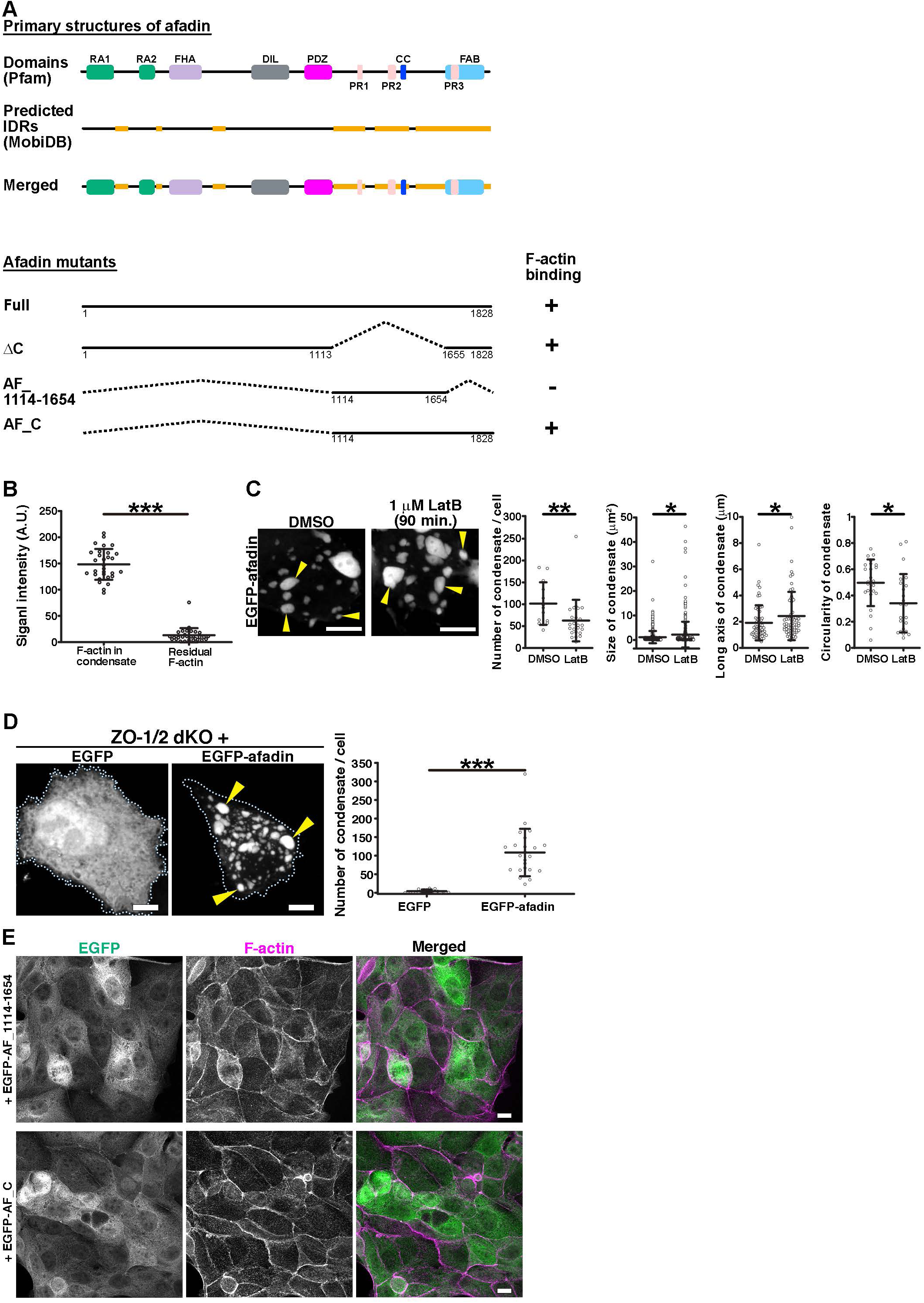
Schematic diagram of afadin IDR mutants and the localization of afadin IDR fragments, related to figure 4. **(A)** Molecular structures of afadin mutants. These afadin fragments were tagged with EGFP at the N-terminus. Predicted IDRs are identified using MobiDB^1^. F-actin-binding (FAB) region was referenced in a previous report^2^. +, positive for the interaction with F-actin; -, negative for the interaction with F-actin. See also Figure S1A. **(B)** Quantification of junctional phalloidin signal in EGFP- or mScarlet- afadin expressing afadin KO cells by the treatment of LatB. Datasets were equal to those in Figure 4F. Whiskers represent mean ± S.D. n = 30 for F-actin in condensate; n = 30 for residual F-actin. Mann-Whitney *U*-test; ***, P ≤ 0.001. See also Figures 4D-G. **(C)** Influence of EGFP-afadin condensates by the treatment of LatB. (Left) Apical views of EGFP-afadin transient expressing HEK293 cells after DMSO or 1 μM LatB treatment for 90 min. Arrowheads indicate EGFP-afadin condensates. Scale bars, 5 μm. (Right) Quantifications of the condensate morphology. Whiskers represent mean ± S.D. Number of condensate/cell; n = 13 for DMSO; n = 24 for 1 μM LatB. Size of condensate; n = 384 for DMSO; n = 353 for 1 μM LatB. Long axis of condensate; n = 64 for DMSO; n = 67 for 1 μM LatB. Circularity of condensate; n = 25 for DMSO; n = 24 for 1 μM LatB. Mann-Whitney *U*-test; **, P ≤ 0.01; *, P ≤ 0.05. **(D)** Transient expression of EGFP or EGFP-afadin in ZO-1/2 dKO MDCK cells. (Left) Apical views of the transfectants cultured for 3 days. Dotted lines represent the cell of interest. Arrowheads indicate condensates. Scale bars, 5 μm. (Right) Quantification of EGFP condensate number of the cells. n = 13 for EGFP; n = 23 for EGFP-afadin. Whiskers represent mean ± S.D. Mann-Whitney *U*-test; ***, P ≤ 0.001. **(E)** Localization of the afadin fragments and F-actin in EGFP-AF_1114-1654 or EGFP-AF_C expressing afadin KO cells. Apical views of the transfectants cultured for 24 h and stained with phalloidin. A representative image is shown of six independent experiments. Scale bars, 10 μm.

**Figure S5.**
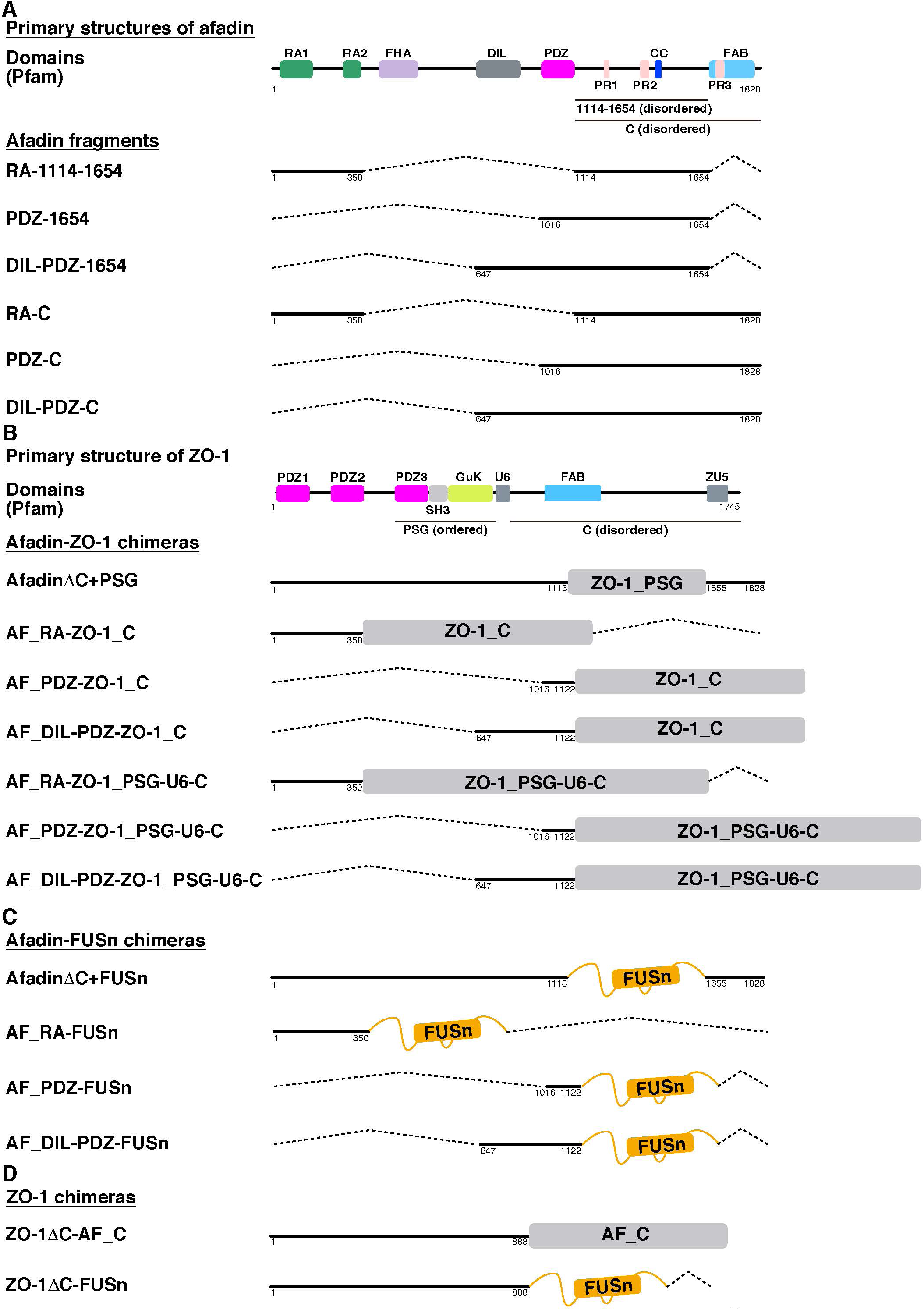
Schematic diagram of systematic afadin extended fragments and ZO-1- Afadin chimeras, related to figure 6&7. **(A)** Molecular structures of afadin extended fragments. These afadin fragments were tagged with EGFP at the N-terminus. See also Figures S1A & S4A. **(B**) Schematic of the reconstructed afadin-ZO-1 chimeras. Ordered ZO-1_PSG requires dimerization of ZO-1 and recruitment of AJ^3^. ZO-1_C contains large IDRs^4-6^. The U6 domain of ZO-1 is the self-inhibited part by folding. Afadin chimeras were tagged with EGFP at the N-terminus. See also Figure S4A. **(C)** Molecular structures of afadin-FUSn chimeras. FUSn is the N- terminal IDR of the RNA-binding protein Fused in Sarcoma (FUS)^7,8^. **(D)** Molecular structures of ZO-1-exogennous IDR attached chimeras. These ZO-1 chimeras were tagged with EGFP at the C-terminus. ZO-1ΔC cannot form condensates in HEK293 cells, and this mutant has a strongly reduced junctional localization in ZO-1/2 dKO MDCK cells^4^.

